# Single Cell transcriptional analysis of *ex vivo* models of cortical and hippocampal development identifies unique longitudinal trends

**DOI:** 10.1101/2022.12.11.519959

**Authors:** Daniel K. Krizay, David B. Goldstein, Michael J. Boland

## Abstract

Postnatal cortical and hippocampal mouse primary neuronal cultures are powerful and widely-used models of neuronal activity and neurological disease. While this model is frequently used to recapitulate what is seen *in vivo*, how the transcriptomic profiles of neuronal networks change over development is not fully understood. We use single-cell transcriptomics to provide a view of neuronal network establishment and maturation. Our data highlight region-specific differences and suggest how cell populations program the transcriptome in these brain regions. We demonstrate that patterns of expression markedly differ between and within neurological diseases, and explore why these differences are found and how well they compare to other models. In particular, we show significant expression differences between genes associated with epilepsy, autism spectrum disorder, and other neurological disorders. Collectively, our study provides novel insights on this popular model of development and disease that will better inform design for drug discovery and therapeutic intervention.

Graphical Abstract
(A) Schematic representing select gene expression progression through neuronal network maturation from human cortical organoids (3- and 6-Month Organoid), newborn mice (P0 Mouse), immature *ex vivo* cortex derived cultures (DIV 3 *ex vivo*), functionally mature *ex vivo* cortex derived cultures (DIV15-31 *ex vivo*), and adult mice (P56 Mouse). Color represents proportion of excitatory neurons with detectable expression for selected representative genes *Mapk10, Igfbp2*, which increase and decrease through network maturation, respectively.
(B) Schematic representing divergent expression patterns between genes associated with epilepsy and ASD through network maturation between the organoids and *ex vivo* cultures shown in (A). Color scales represent the change in the percentile, in respect to all genes, of the proportion of excitatory neurons with detectable expression.

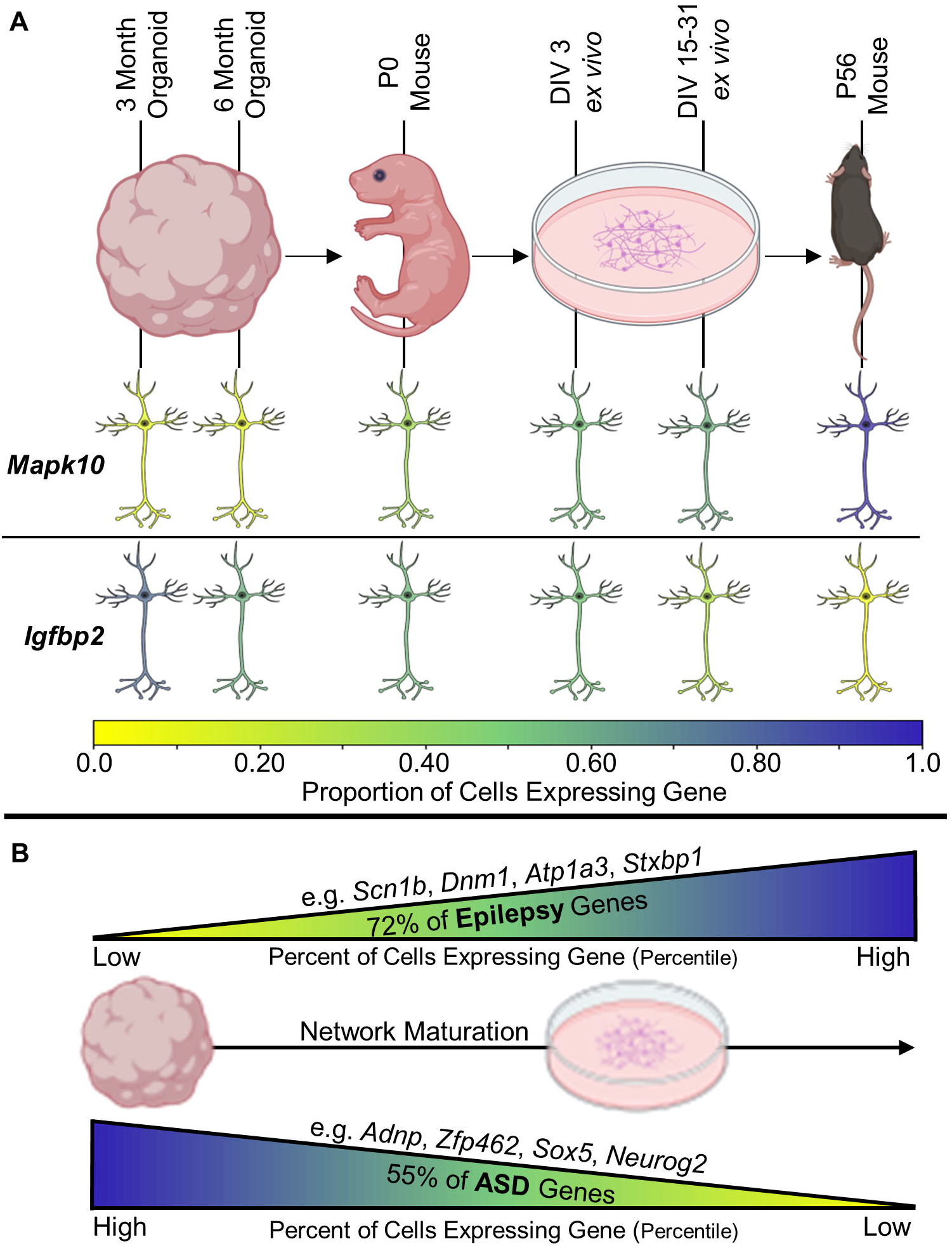

## Introduction

Transcriptomic dynamics during developing embryonic and postnatal brains are vital for a better understanding on how (epi)genetic programming underlies the complex processes of proliferation, neuronal maturation, network and circuit formation, and ultimately electrophysiological behavior and disease. Models of neurological diseases are used to investigate these questions, including mouse models, primary mouse neuronal cultures, and human induced pluripotent stem cell (hiPSC)-derived brain organoids and two-dimensional cultures[1–4]. Mice are often used because of their large litters, short lifespans, and ease of genetic manipulation[5], but sometimes fail to recapitulate prominent and complex phenotypes of human diseases due to the inherent differences in the spatiotemporal dynamics of brain development, architecture, and function between the mouse and human brains [6–9]. HiPSC-derived brain organoids are commonly used to model neurological diseases[3, 10–14] such as cortical malformations of development[15–17], ASD[18–21], and genetic epilepsy[22–24], and provide the benefit of being derived from human cells while allowing for a structural architecture that resembles that which is seen *in vivo*[25, 26]. However, organoids represent an artificial construction of the brain that lacks many essential features that cannot yet be accurately and consistently modeled[27, 28], suffer from issues in reproducibility[29–31], and exhibit the protracted maturation of human development, which requires months to achieve an organoid with relative immaturity[25, 32, 33]. In comparison, 2D hiPSC cultures provide a cheaper and more reproducible human-derived model[34, 35], but lack structural 3D complexity. In addition, both hiPSC models have multiple differentiation protocols which preferentially enrich for specific neuron populations, leading to biologically inconsistent and inaccurate ratios of excitatory and inhibitory neurons[36–39], and the complete absence of some important neuronal subtypes[40–43].

Mouse primary *ex* vivo cultures also lack the structural complexity of 3D models, but benefit from their relative ease of use, low costs, high reproducibility, and relative functional maturity[44–46]. These merits lead to the frequent use of this model for neurological and developmental disorders [47–49], neurophysiology[50–52], and drug screening[44, 47, 53, 54]. While these models can be transcriptionally informative, like in their *in vivo* counterparts, the expression of many genes is temporally dynamic[55, 56]. This is especially true during early postnatal periods when neurodevelopment is very active[57, 58]. The extent and specifics of the dynamic temporal gene expression in these postnatal cultures over network development is not well characterized, and is crucial for experimental design when using these cultures, or deciding which model to use. This is of increased importance when looking at genes that are known to be causal for, or associated with, neurological disorders[59–67].

To address this problem, we generated a longitudinal transcriptomic profile of postnatal developing mouse primary cortical and hippocampal cultures. We uncovered unique developmental transcriptomic profiles for individual genes, disease gene subclasses, and biological processes. Single cell profiling uncovered the distinct developmental trajectories of cell populations that would be lost or diminished in bulk RNA sequencing. Understanding these cell-type specific transcriptomic differences is especially important in the research of genetic neurological disorders, as it is known that transcriptional variations in specific genes of interest lead to functional abnormalities in specific cell subtypes [68, 69]. Specifically, we discovered cell population-specific divergent transcriptomic profiles between genes associated with neurological diseases, focusing on epilepsy and autism spectrum disorder.

We also compared the data from our *ex vivo* system, to transcriptomic data collected from *in vivo* neonatal and adult mouse brains and human cortical organoids. We discovered that while the expression profiles between the systems are relatively correlated, system-wide differences as well as gene-specific differences are present, finding an enrichment for immature biological pathways in 6-month organoids as compared to neonatal mice. We discuss this relative immaturity of cortical organoids, and how this impacts the choice of which model system to use when researching genes of interest, specifically those associated with neurological disorders. This paper highlights the importance of the generation and consideration of system-specific transcriptomic datasets when looking into a gene, disease, or biological process of interest, and serves as a vital resource for researchers while providing a more complete and granular analysis of ex vivo postnatal development.

## Results

### Longitudinal transcriptomic analyses of region-specific *ex vivo* neuronal network maturation

Based on longitudinal activity data collected from primary cortical and hippocampal neuronal cultures using multi-electrode array (MEA), we examined neuronal activity during network establishment to select time developmental time points of interest for transcriptomic interrogation [47-49] (Figure 1A). We collected the cortices and hippocampi from twelve postnatal day zero (P0) male C57BL/6J mice from three separate litters and plated a portion of cells for MEA, immunocytochemistry (ICC) and single-cell RNA-sequencing (scRNA-seq). We used MEA recordings to verify that the cultures became functionally active in an expected manner indicating healthy network establishment (Figure 1A). On Days *in vitro* (DIV)3, DIV9, DIV15, DIV23, and DIV31, cells were collected and submitted for sequencing, and parallel cultures on coverslips were fixed and stained for cell-type specific markers (Figure 1B). The ICC images verify that the cultures were healthy and contain a variety of expected cell types. (Figure S1).

**Figure 1.**
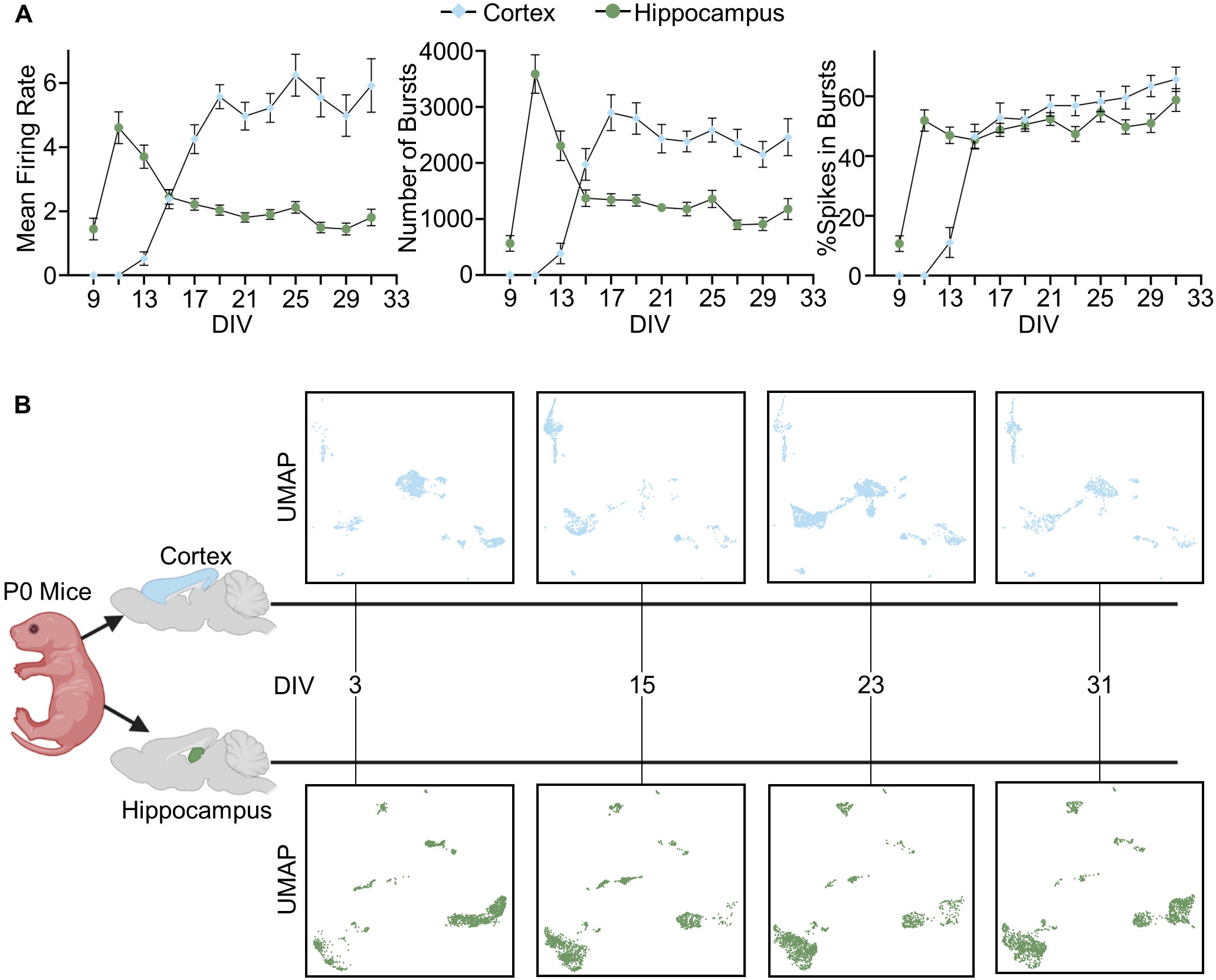
*Ex vivo* neuronal network activity informs experimental design (A) *Ex vivo* cortical (blue) and hippocampal (green) primary cultures recorded every other day on multi-electrode arrays (MEA) from DIV9-DIV31. Mean firing rate relative to the number of active electrodes, total number of bursts, and the percent of spikes in bursts displayed from left to right. Cortical cultures, n = 24. Hippocampal cultures, n = 24. Error bars indicate SEM. (B) Schematic of experiment design. UMAP plots of scRNA-seq data for each of the four timepoints (DIV3, DIV15, DIV23, DIV31) analyzed from mouse cortical (blue) and hippocampal (green) primary cultures.

Raw sequencing reads (Table S1) were QC’d and analyzed to filter against dead and damaged cells and artifactual libraries derived from cell doublets and extracellular reads. From this data, unsupervised clustering using Seurat v4[70–72] was performed (Figure S2). DIV9 data from both the cortical and hippocampal cultures were excluded from downstream analysis as the median genes per cell were too low to provide valuable information from this time point (Table S1). Following QC and excluding the DIV9 samples, our data set is comprised of 5,802 cortical cells and 6,774 hippocampal cells in total (Table S2). Using canonical cell-type specific markers (Figure 2G,H) [48, 73–76], we curated and combined clusters into broad cell populations (Figure 2A:F). We identified the expected cell populations (excitatory neurons, inhibitory neurons, astrocytes, oligodendrocytes/OPC (OPC), smooth muscle cells (SMC), and Cajal-Retzius cells) (Figure 2A, D; Table S2). While we are combining our cortical neurons into broad classes, it is important to note that we do see layer specific populations, including upper-layer neuronal marker positive clusters (*Satb2* and *Cux2*), and deep-layer neuronal marker positive clusters (*Bcl11b* and *Tbr1*). For the hippocampal neurons, we also see distinct hippocampal region-specific clusters, including DG neuronal marker positive clusters (*Prox1*) and CA1 neuronal marker positive clusters (*Wfs1* and *Mpped1*). We performed downstream analysis on excitatory and inhibitory neurons and astrocytes due their involvement in neurodevelopmental disorders and their utility for drug screening.

**Figure 2.**
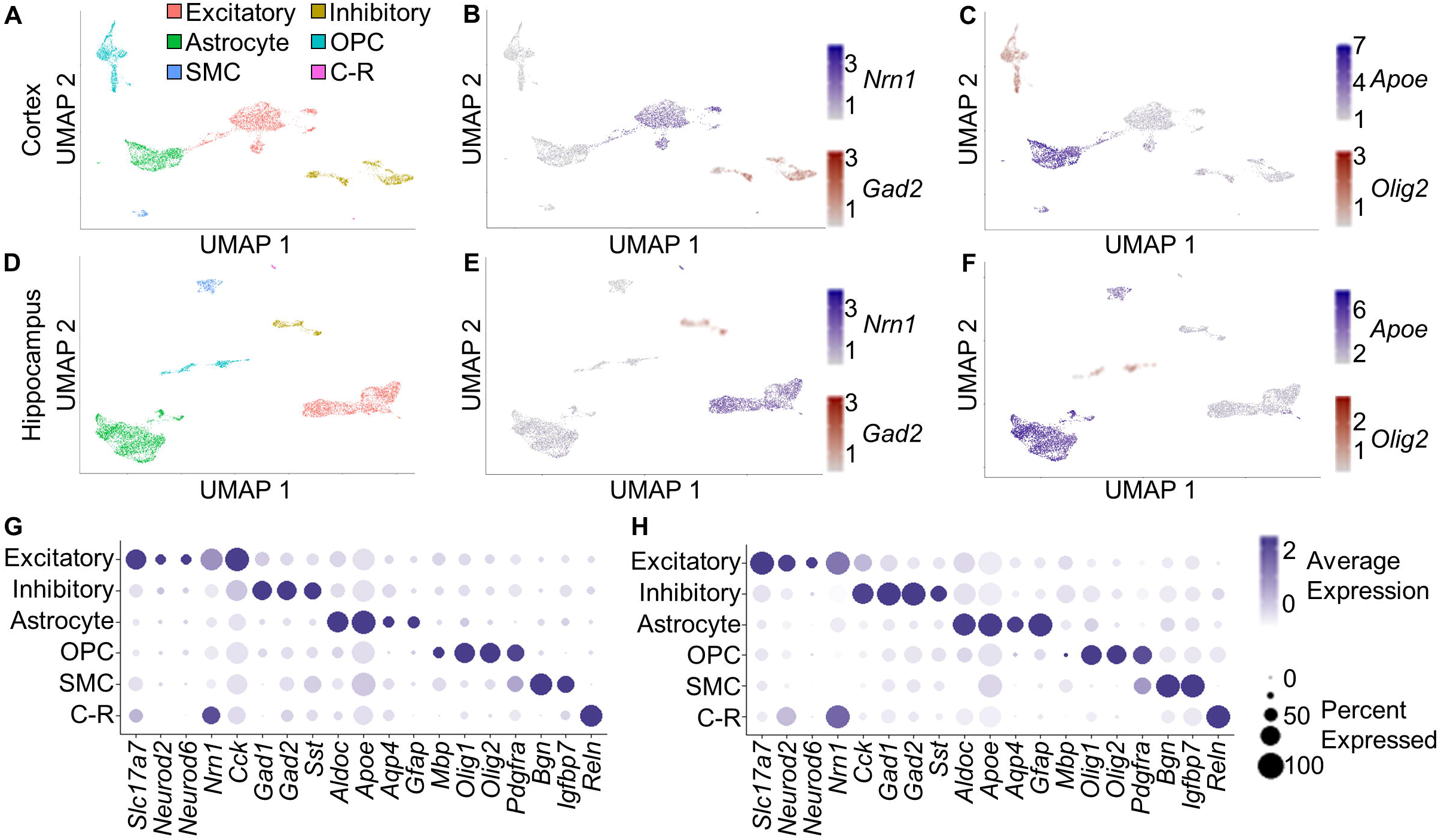
Annotation of wildtype mouse cortex and hippocampus derived *ex vivo* cultures. (A) UMAP plot of post-QC scRNA-seq data across all time points from 5,802 mouse cortical cells collected across 12 mice. Color represents broad cell population clusters. (B) Expression levels of *Nrn1* and *Gad2* for cells in (A) represented with respective color intensities. (C) Expression levels of *Apoe* and *Olig2* for cells in (A) represented with respective color intensities. (D) UMAP plot of post-QC scRNA-seq data across all time points from 6,774 mouse hippocampal cells from 12 mice. Color represents broad cell population clusters. (E and F) Same as B and C for hippocampal cultures (D). (G) Dot plot displaying the expression of select cell-type-specific marker genes (columns) in the broad cell populations found in (A) and (D) for cortex. Dot size represents percentage of cells expressing a marker gene, and color intensity represents relative average expression of a marker gene. (H) Same as G for hippocampus. Excitatory, Excitatory Neurons; Inhibitory, Inhibitory Neurons; OPC, Oligodendrocytes and Oligodendrocyte Precursor Cells (OPC); SMC, Smooth Muscle Cells (SMC); C-R, Cajal-Retzius Neurons.

### Enriched biological processes support proper network development

In addition to the average expression level for each gene, we calculated the percentage of cells expressing each gene within each of the cell classes. As average expression of each gene is heavily driven by the percentage of cells expressing a gene, we also looked at the average expression of each gene with the cells not expressing each gene omitted from the calculation (Table S3). To determine which biological processes were enriched in both immature and mature cultures, we found the top 500 genes with the greatest difference between DIV3 and DIV31 for these calculations and performed gene ontology (GO) analyses[77–79] (Figure 3A; Table S4). In both excitatory and inhibitory neuron populations for both the cortical and hippocampal cultures, genes more highly expressed in DIV3 neurons than DIV31 neurons are enriched for pathways related to immature and developing neurons, including nervous system development, neurogenesis, and neuron differentiation.

**Figure 3.**
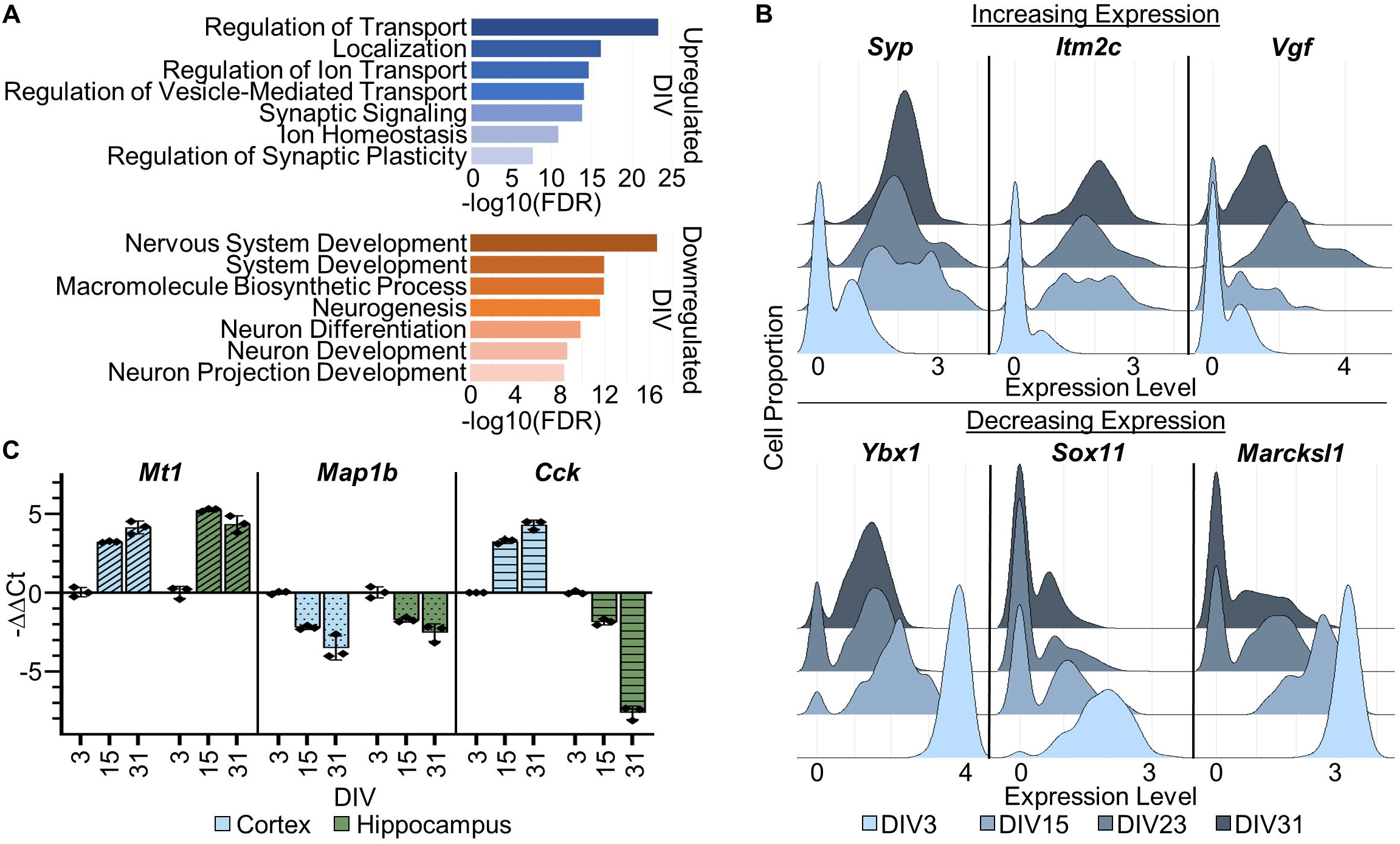
Temporal analysis of *ex vivo* cultures demonstrates biologically consistent transcriptomic trends (A) GO analysis of 500 genes with largest difference in average expression between DIV03 and DIV31 (Blue plot: DIV03 < DIV31, Orange plot: DIV03 > DIV31). Bar represents –log10 value of FDR adjusted p-value for each of the representative enriched biological processes. (B) Ridgeline plot of the cortex excitatory neuron population for three representative genes with gene expression positively correlated with increasing culture age (Top) and negatively correlating with increasing culture age (Bottom). Color represents cell culture age. (C) Bar plot with individual datapoints representing RT-qPCR data for three representative genes, one with gene expression positively correlated with increasing culture age in both *ex vivo* cultures (*Mt1*), one negatively correlated with increasing culture age in both *ex vivo* cultures (*Map1b*), and one positively correlated with increasing culture age in cortical (blue) derived cultures, and negatively correlated with increasing culture age in hippocampal (green) cultures (*Cck*). Bars represent −ΔΔCt values of each gene for DIV3, 15, and 31 relative to DIV3 for each group. Each datapoint represents the average of two technical replicates, and error bars represent the standard deviation of the three biological replicates. Gnb1 was used as the reference control gene for all samples.

For the cortical excitatory neuron population, genes more highly expressed in DIV31 neurons than DIV3 neurons are enriched for pathways related to mature and active neuronal cultures, including regulation of ion transport, synaptic signaling, and regulation of synaptic plasticity. Cortical inhibitory neurons and hippocampal excitatory neurons are enriched for similar transport and localization pathways as cortical excitatory, but are not as significantly enriched for synaptic/neuronal firing pathways. However, hippocampal inhibitory neurons possess immature pathways as seen in DIV3 profiles relative to DIV31. This indicates that inhibitory neurons in the hippocampus are not as mature as cortical inhibitory neurons in these cultures. In addition, the Pearson correlation of gene expression levels for all genes between the four timepoints (for both cortical and hippocampal cultures) shows that DIV3 was less correlated to the other timepoints than they were with each other, which is likely due to DIV3 having the most dynamic developmental potential of the four time points as it is still early in network establishment relative to later time points (Figure S3). As expected, the excitatory and inhibitory neuron classes were more correlated with each other than with astrocytes.

We then examined specific genes involved in well-characterized and temporally sensitive pathways. These include genes involved in synaptic activity, negative regulation of development, and synaptic plasticity (*Syp, Itm2c, Vgf*) and genes involved in development and differentiation (*Ybx1, Sox11, Marcksl1*) to verify that individual genes have expected expression changes (Figure 3B; Figure S4A). We found that for all genes analyzed, the expected gene expression trends were found, supporting the validity of these datasets as a model of network establishment.

We screened our scRNA-seq data to find genes with significantly large fold change differences between DIV3 and DIV31 (Log_2_ fold change of >0.75 or <-0.75 and an FDR adjusted p-value of <0.01) in both the cortex and hippocampus derived cultures. From this subset of genes, representative genes *Mt1, Map1b*, and *Cck* were selected, as these genes, in respect to culture age, represented those with increased expression for both cortex and hippocampus derived cultures, decreased expression for both cortex and hippocampus derived cultures, and increased expression for cortex derived cultures but decreased expression for hippocampus derived cultures, respectively (Table S5). The longitudinal gene expression profiles of these three genes were verified via RT-qPCR in an independent set of cultured neuronal networks (n = 3, Figure 3C) using *Gnb1* as the internal control gene as it exhibits highly stable expression over time between cell populations from both brain regions (see below).

### Introduction of pseudotime analysis increases temporal granularity

In order to increase the granularity of cell maturation and age beyond only looking at a given gene’s expression by DIV, we generated diffusion maps for each of the three major cell populations within both the cortical and hippocampal datasets [80]. From these maps, we can place cells along an expected differentiation and maturation trajectory, assigning each cell a diffusion pseudotime value that is representative of its relative age and maturation in respect to all cells in the dataset. We then determined the pseudotime correlation value (PCV) of each gene in our cell populations of interest by calculating the correlation between each gene’s expression and the assigned pseudotime value for each cell (Table S3). With this value, we determined how each gene’s expression changes over cell development through a more granular approach. A positive PCV represents a gene with increasing expression over the maturation of the cell population, while a negative PCV represents a gene with decreasing expression over time.

We determined a correlation between the PCV and the DIV gene expression correlation calculated above to determine how accurate this method of maturation trajectory worked with these postnatal datasets. There was strong positive correlation in all cell populations tested, ranging between 0.67 and 0.99 (Figure 4A). We performed GO enrichment analysis for each cell population to determine which biological processes were enriched in the genes with the highest and lowest PCV (top 500 genes) (Figure 4B; Table S6). For cortical and hippocampal excitatory neurons, genes with the lowest PCV are enriched for developmental and differentiation pathways. For cortical excitatory and inhibitory neurons, and hippocampal inhibitory neurons, genes with the highest PCV are enriched for pathways involved in synaptic activity and transport. Genes with detectable expression related to the representative GO pathways, synaptic signaling and peptide biosynthetic process show a significant trend of upregulation and downregulation, respectively, of gene expression from DIV3 to DIV31 in excitatory neurons. (Figure S5; Table S7). Genes with the lowest PCV in cortical and hippocampal inhibitory neurons were enriched for metabolic, biosynthetic, and translational pathways, unlike what was seen in the excitatory neurons. In hippocampal excitatory neurons, the genes with high PCV are not enriched for pathways associated with synaptic activity, and instead are only enriched for ion transport and localization pathways. One key note is that compared to biological processes found analyzing gene expression in DIV3 vs. DIV31 timepoints, these PCV results are more biologically consistent with what we would expect to see, giving further validity to this diffusion pseudotime approach.

**Figure 4.**
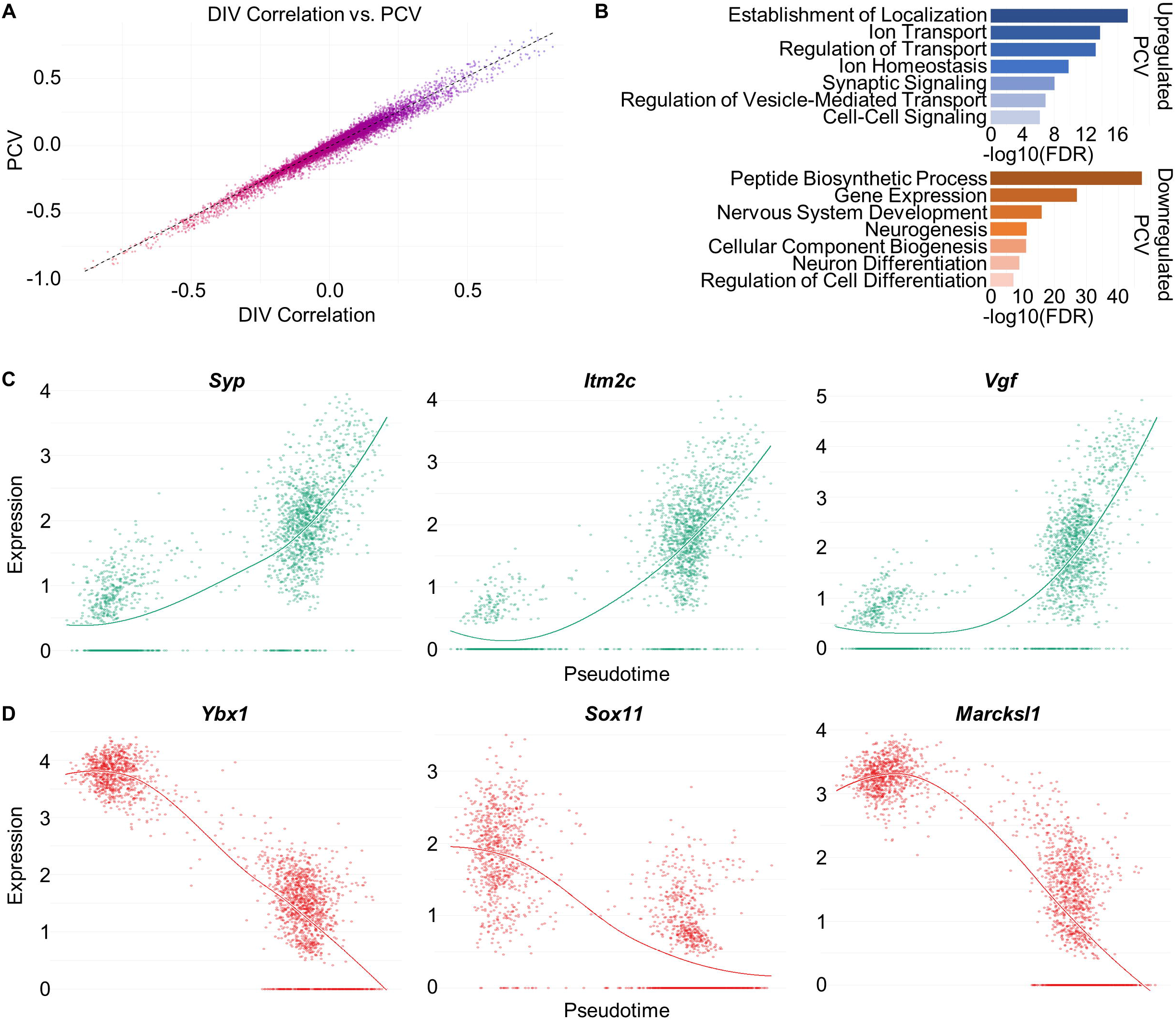
Pseudotime analysis of *ex vivo* cultures shows biologically consistent transcriptomic trends (A) Scatter plot representing the correlation between the Pearson correlation of gene expression to culture age (“DIV Correlation” on x-axis) and the correlation of gene expression to pseudotime (“PCV” on y-axis) for the cortex excitatory neuron population. Color gradient represents correlation value along the first principal component of the plot. Dotted line represents linear regression line for the data. (B) GO analysis for the top 500 genes with highest (Blue) and lowest (Orange) pseudotime correlation values (PCV). Bar represents –log10 value of FDR adjusted p-value for each of the representative enriched biological processes. (C) Scatter plots comparing gene expression and pseudotime for the three genes in (Fig. 3B) that are positively correlated with culture age for each cell in the cortical excitatory neuron population. The line on each plot represents a smooth local regression of the data. (D) Same as (C) for the three representative genes shown in (Fig. 3B) that are negatively correlated with culture age.

To determine differences between cell populations within brain regions, we took the 500 genes with the greatest difference in PCV with a positive PCV in one population and negative PCV in another and determined whether these genes are significantly enriched in certain biological pathways. In both cortical and hippocampal cultures, genes with a positive PCV in inhibitory neurons but negative PCV in excitatory neurons are significantly upregulated in processes related to nervous system development, neurogenesis, and differentiation (Figure S4B), suggesting that inhibitory neurons have a slower developmental trajectory in our *ex vivo* cultures than excitatory neurons, a finding that agrees with the protracted maturation of inhibitory neurons *in vivo*[81–83].

In addition to looking at the macro trends, we can once again look at specific representative genes for synaptic activity and development to verify expected positive or negative PCVs in our system. Looking at the same genes as before, this representative expression pattern is replicated in all examples (Figure 4C,D; Figure S4C,D). We also see a highly positive PCV with *Slc12a5* (KCC2), which is known to have increased expression over time, and contribute to the GABA functional switch. When taken together, these data highlight the strength and validity of diffusion pseudotime analysis for analyzing longitudinal transcriptomic data in this model.

### Cortical and hippocampal derived cultures demonstrate unique cell type-specific development

We compared the expression patterns between the development of cortical and hippocampal cultures. Overall, the PCV correlation between the brain regions for both the excitatory and inhibitory neuron populations was high (0.80), while displaying much lower correlation for the astrocyte population (0.65). The large differences in functional activity of our cortical and hippocampal cultures (Figure 1) could be attributed to transcriptomic differences between brain regions. Therefore, we determined which genes show diverging PCV patterns between the cortical and hippocampal cultures. In excitatory neurons, we identified a subset of genes with increasing expression in the cortex and decreasing expression in the hippocampus (e.g. *Cck, Snca*) as well as genes with decreasing expression in the cortex and increasing expression in the hippocampus (e.g. *Slc25a5, Eif1b*) (Figure 5A).

**Figure 5.**
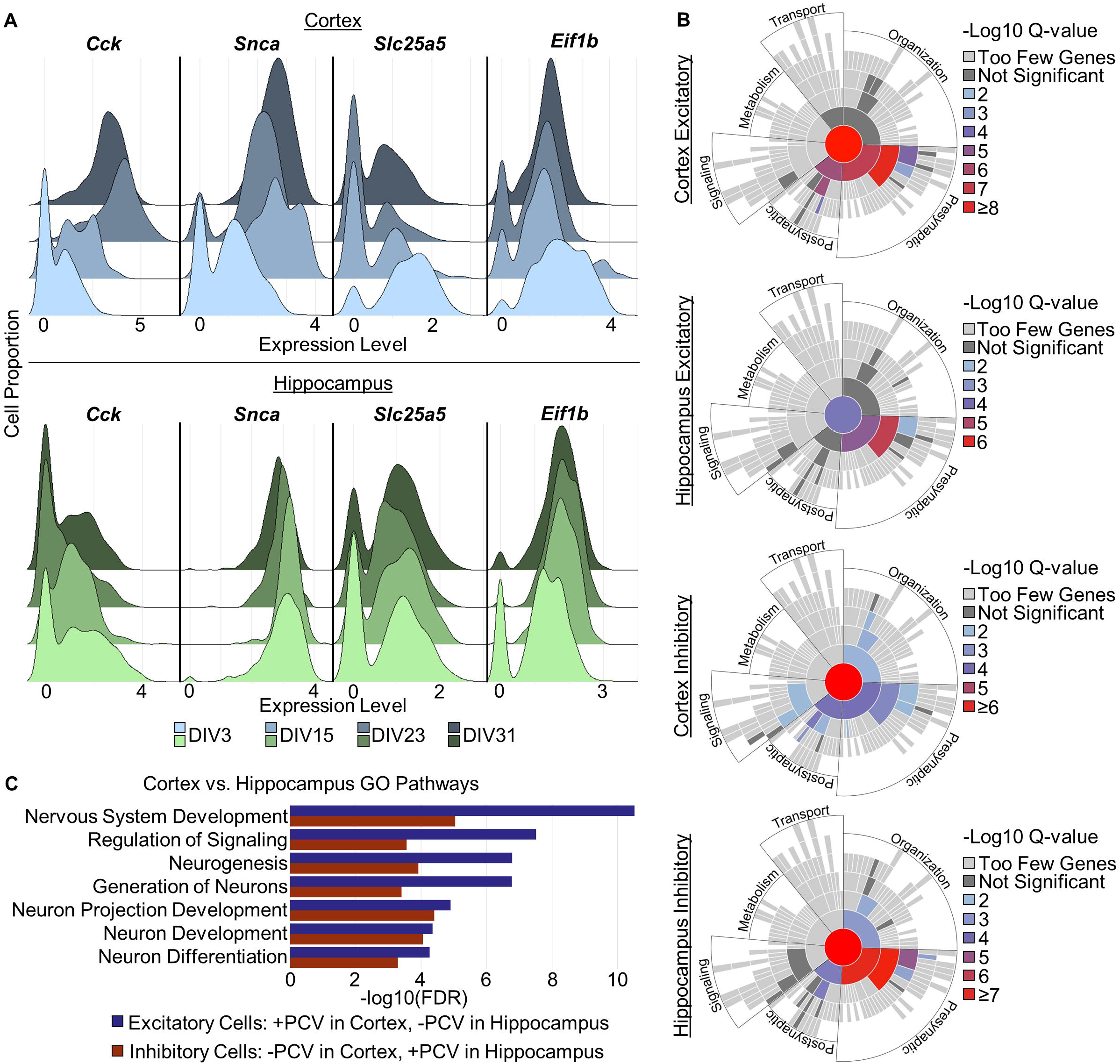
Brain region-specific cultures have unique longitudinal transcriptomic profiles (A) Ridgeline plots for four representative genes with gene expression differences between the cortex excitatory neuron population (top) and the hippocampus excitatory neuron population (bottom). Color represents cell culture age. (B) SynGO plots for top 500 genes with largest PCVs for cortex excitatory neurons (1st), hippocampus excitatory neurons (2nd), cortex inhibitory neurons (3rd), and hippocampus inhibitory neurons (4th). Colors represent –log10 Q-values for enriched synaptic processes. Top-level synaptic processes are labeled. (C) GO analysis of top 500 gene sets with the greatest difference in PCV where a gene has a positive PCV in one brain region and a negative PCV in the other. Blue bars represent the gene set with positive PCVs in cortical excitatory neurons and negative PCVs in hippocampal excitatory neurons. Red bars represent the gene set with negative PCVs in cortical inhibitory neurons and positive PCVs in hippocampal inhibitory neurons.

Synaptic Gene Ontology (SynGO) analysis was performed using stringent evidence filters for the 500 genes with the highest and lowest PCV values in the excitatory and inhibitory neuron populations (Figure 5B). For both neuronal populations, the genes with the lowest PCVs had no significant enrichment for synaptic processes. The genes with the highest PCVs in cortical excitatory neurons showed significant enrichment in presynaptic and postsynaptic processes, including synaptic vesicle exocytosis and neurotransmitter receptor localization, respectively. This differs from the hippocampal excitatory neurons, where only presynaptic processes are enriched. The top PCV genes in cortical inhibitory neurons showed significant enrichment in presynaptic, postsynaptic, synapse organization, and signaling processes. In comparison, hippocampal inhibitory neurons showed much stronger enrichment in presynaptic processes and no enrichment in signaling processes.

General GO analysis was also performed to compare the cortical and hippocampal cultures. Genes with a positive PCV in cortical excitatory neurons and a negative PCV in hippocampal excitatory neurons are enriched for biological processes associated with immature neurons, like nervous system development, cell differentiation, and neurogenesis (e.g. *Cck, Epha5, Cnr1*) (Figure 5C). Interestingly, this trend is reversed for inhibitory neurons, where genes with a positive PCV in hippocampal cells but a negative PCV in cortical cells are enriched for these same biological processes (e.g. *Dpysl2, Elavl3, Gsk3b*). We interpret this, in combination with the SynGO data, to mean that excitatory neurons develop faster in the hippocampus than in the cortex, whereas hippocampal inhibitory neurons exhibit more protracted development relative to cortical interneurons. Interestingly, the genes that make up these sets largely do not overlap but the processes do. Together, these findings may partially explain the differences in MEA activity profiles between the two regions where the hippocampus derived cultures exhibit increased firing early in development (DIV11 relative to cortex), before decreasing and subsequent plateau, while cortex derived cultures show a low amount of early activity that gradually increases before plateauing around DIV19 (Figure 1A).

### Genes with stable expression suggest better alternatives to customary reference genes

While looking at genes with temporal expression differences is important to better understand neurodevelopment, we also identified ubiquitously expressed genes that show stable expression throughout postnatal development, as genes such as these are often used as internal controls to normalize transcriptomic data. We first interrogated the expression stability of 22 widely-used “housekeeping” genes [84, 85] through the development of these cultures (Table S8). We defined a good housekeeping gene as one that is highly expressed in all cell types (expressed in >50% of cells within a given population) while keeping stable expression throughout the development of the cultures (i.e. PCV close to 0). Of the 22 genes investigated, only 10 were expressed in greater than 50% of all cells, and of those 10, all have at least one cell population with an absolute PCV >0.2, with the lowest being *Ubc* (PCV 0.27) and the highest being *Rplp0* (PCV 0.79). As none of these genes fulfilled our definition of a housekeeping gene, we mined our data to discover better candidate housekeeping genes. Using the same cell proportions (>50%) and PCV filters (absolute value <0.2), we discovered 13 genes (Table S8; Figure S6) that exhibited stable temporal expression in a majority of the cells within each cell type in each brain region (e.g. *Gnb1, Ndufb5, Maged1, Psenen*). These genes have roles in biological processes including mitochondrial organization, cellular localization, intracellular transport, and protein localization. Our data suggest these genes are better housekeeping genes than any of the genes commonly used (e.g. *Gapdh, Actb, Rpl13*).

### Genes associated with neurological disorders show unique temporal transcriptomic trends

Next we curated and investigated the gene expression patterns and cell specificity of 94 genes associated with genetic epilepsy (compiled and curated from [59, 60]), 227 Autism Spectrum Disorder (ASD) genes (SFARI Category 1 and SPARK genes[61, 62, 86, 87]), 32 schizophrenia genes [63, 88], 16 dystonia genes [64, 65], 84 genes associated with Alzheimer’s Disease [66], 55 Parkinson’s disease genes from the Gene4PD database [67], 57 Obesity genes [89], and 87 Coronary Artery Disease (CAD) genes [90, 91] (Table S9).

We calculated the percentile of the percentage of cells expressing each of these disease genes for each cell population in both the cortex and hippocampus as compared to the group of all detected genes (Table S10). We found that on average, the neurological disorder genes are expressed in a relatively large percentage of the cells (76^th^ percentile), ranging from 74^th^ −80^th^ percentiles. The obesity and CAD genes, however, were expressed in a lower percentage of cells (68^th^ and 64^th^ percentile respectively). For reference, the set of all genes expressed in our cultures are, on average, expressed in 15.7% and 17.8% of cells in the cortex and hippocampus respectively. All of this together suggests that neurological disease genes are more highly expressed in our cultures than non-neurological disease genes, and disease genes, as a whole, are expressed in more cells than the average of all genes.

We then compared the average PCV for each disease gene in each cortical and hippocampal cell population to the average PCV of the entire gene dataset in these cell types. The average PCV for the epilepsy genes in cortical and hippocampal excitatory and inhibitory neurons was determined to be in the 80^th^ to 89^th^ percentile of all genes (Figure 6A). The top 25% of epilepsy genes by PCV were enriched for ion transport and trans-synaptic signaling, while the bottom 25% were enriched for axon development and developmental processes.

**Figure 6.**
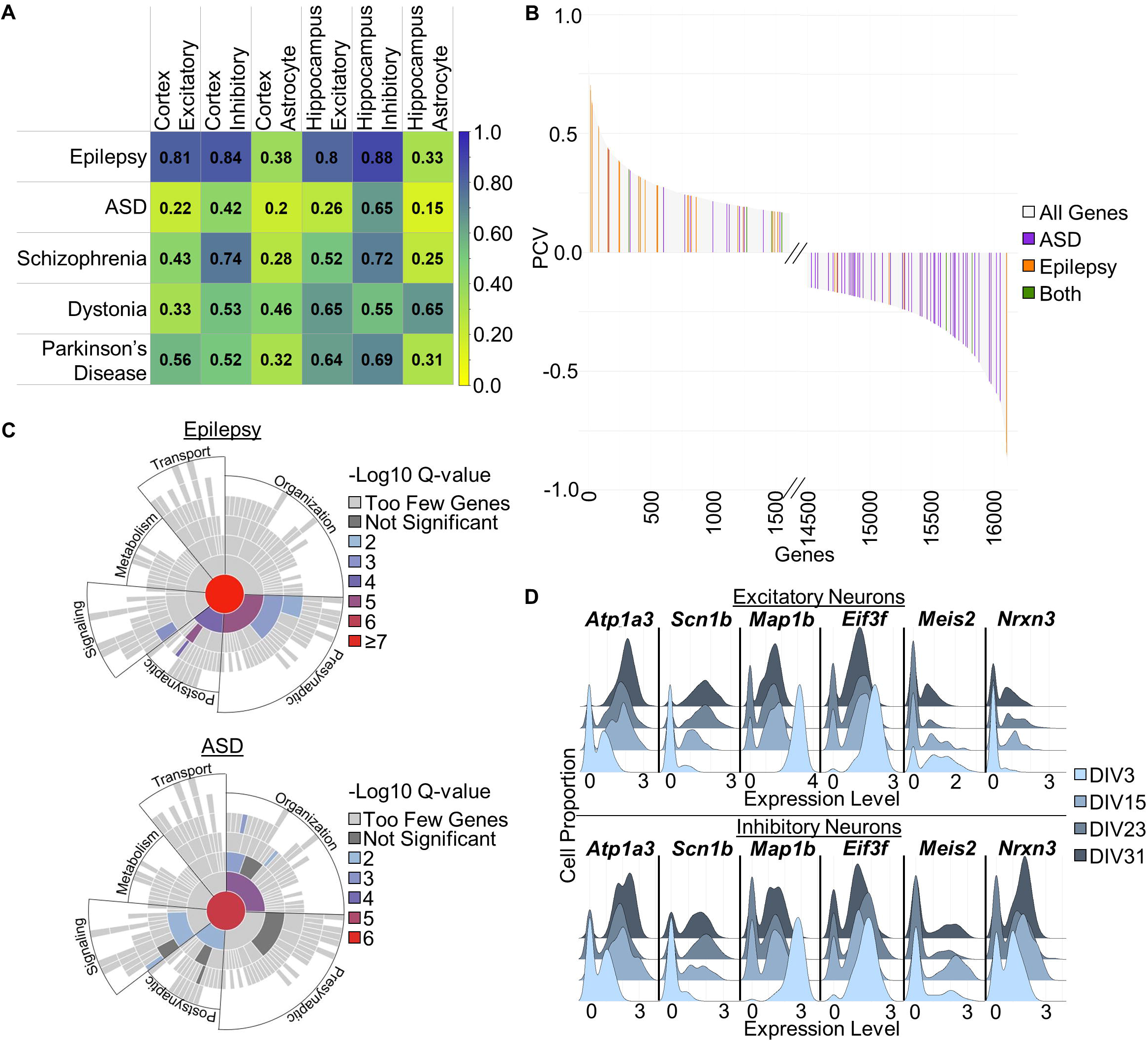
Unique longitudinal transcriptomic trends found between neurological diseases (A) Table displaying the percentile from, 0 to 1, for the average PCV for all genes in the neurological diseases analyzed (rows), in each of the six major cell populations tested (columns). Yellow represents low average PCV and blue represents high average PCV. (B) Bar plot of top and bottom 10% of all genes plotted from highest to lowest PCV along x-axis for cortex excitatory neurons. ASD genes, epilepsy genes, and genes associated with both ASD and epilepsy are represented with purple, orange, and green bars respectively. (C) SynGO plots for genes associated with epilepsy (top) and ASD (bottom). Disease gene lists can be found in Table S8. Colors represent –log10 Q-values for enriched synaptic processes. Top-level synaptic processes are labeled. (D) Ridgeline Plots for cortex excitatory neuron (top) and cortex inhibitory neuron (bottom) populations for six representative genes. *Atp1a3* and *Scn1b* represent genes with positive PCVs in both populations, *Map1b* and *Eif3f* represent genes with negative PCVs in both populations, and *Meis2* and *Nrxn3* represent genes with differing PCVs between both populations. Color represents cell culture age.

For ASD genes, the average PCV for cortical and hippocampal excitatory neurons were in the 22^nd^ and 26^th^ percentile respectively, with the bottom 25% being enriched for biological pathways including development and differentiation. Interestingly, this trend of low percentile ASD genes was not seen in the inhibitory neurons. When genes that were classified as both Epilepsy and ASD genes are removed from the analysis, this divergent trend further strengthened for both.

For schizophrenia associated genes in inhibitory neurons in the cortex and hippocampus, the average PCV was also high (77^th^ and 74^th^ percentile respectively). This trend is not seen in the excitatory neurons for both brain regions. The results for the other disease classes can be found in Figure 6A and Table S10. We plotted the PCV for each gene in the cortical excitatory neuron population from highest to lowest, with the Epilepsy and ASD genes highlighted (Figure S7). The top 10^th^ and bottom 10^th^ percentile of PCV are highlighted to show the differing distribution of epilepsy and ASD genes in more detail (Figure 6B).

SynGO analysis was performed on each of the disease classes with stringent evidence filters to determine how each disease gene set is uniquely represented in synaptic processes (Figure 6C; Figure S8). After removing overlapping genes, genes associated with genetic epilepsy are enriched for presynaptic and postsynaptic processes including synaptic vesicle exocytosis and regulation of postsynaptic membrane potential, while ASD genes are enriched for synapse organization and trans-synaptic signaling processes including regulation of postsynapse organization and modulation of chemical synaptic transmission. 39% of epilepsy genes were mapped to unique SynGO genes, as compared to only 22% of ASD genes.

In addition to understanding trends of disease classes, it is also important to understand which, and how, individual genes within these classes are driving the trends we have found within and between the brain regions and cell populations. In both excitatory and inhibitory neurons, very high PCV disease genes (e.g. *Atp1a3, Scn1b*) and very low PCV disease genes (e.g. *Map1b, Eif3f*) are partially causal for these disease trends. While genes with divergent expression patterns (e.g. *Meis2, Nrxn3*) between the neuronal subtypes are partially causal for the differences seen between cell populations (Figure 6D). We also looked at which genes have no detectable expression in our cell populations of interest (Table S11). Of the 508 disease associated genes we analyzed, we were unable to detect expression in 26 genes, including but not limited to epilepsy genes *Casr, Chrna2*, and *Gpr98*, and ASD genes *Hectd4, Kmt5b*, and *Tek*. 48 of our disease genes show no detectable expression in at least one of our cell populations of interest. Looking at it deeper, we also found genes with no detectable expression in the cortex but expression detected in the hippocampus including *Pax5, Agtr1a*, and Mug1, with the reverse trend found in *Cd33, Cpa6*, and *Pah*. Epha1 had no expression in our neuronal cell populations, while *Adra2b* had no detectable expression in only excitatory neurons. In contrast, some genes including but not limited to *Echdc3, Hydin*, and *Cd33* have no detectable expression in inhibitory neurons, but do show expression in excitatory neurons. More detailed expression patterns can be found for these 48 genes in Table S11.

### Analysis of developmental gene expression between models, species, and maturity

In an effort to determine how different models of cortical development compare, we analyzed scRNA-seq data from 3 and 6-month human cortical organoids, P0 mice dissociated cortices, and 8-week-old adult mice dissociated cortices. We analyzed expression patterns within and between different developmental models, ages, and species to gather a more complete understanding of each model’s strengths and weaknesses. Human cortical organoid scRNA-seq data was acquired from 3-month and 6-month-old dorsally patterned organoids[29].P0 mouse cortical scRNA-seq data was acquired from C57Bl/6NJ dissociated cortices[48]. Adult (8-week-old) mouse cortical scRNA-seq data was acquired from C57BL/6J dissociated cortices[74]. Detailed information on data origin and data processing can be found in ‘Methods’ section. The data from these three new models were QC’d, processed, and analyzed using the same methods as done with our *ex vivo* datasets, and were clustered into major broad cell populations (Figure S9A:C). From these, we performed further analysis on the same three major cell populations of excitatory neurons, inhibitory neurons, and astrocytes (Tables S12, S13, S14).

We took the set of all genes with detectable expression in any of the datasets, and analyzed how average gene expression for all genes correlates between each of the five developmental mouse timepoints we investigated (P0, DIV3, DIV15, DIV23, DIV31) (Figure 7A). Within each of the three major cell populations analyzed, the correlation between timepoints was high, ranging from 0.84-0.98. Strikingly, the P0 dataset correlated most with DIV3 (0.96-0.97) and least with DIV23 and DIV31 (0.84-0.89), while DIV31 correlated most with DIV23 and DIV15 (0.96-0.98) and least with P0 (0.85-0.89), showing a consistent gradient of expression similarities as the biological age of these mouse models of cortical development increase.

**Figure 7.**
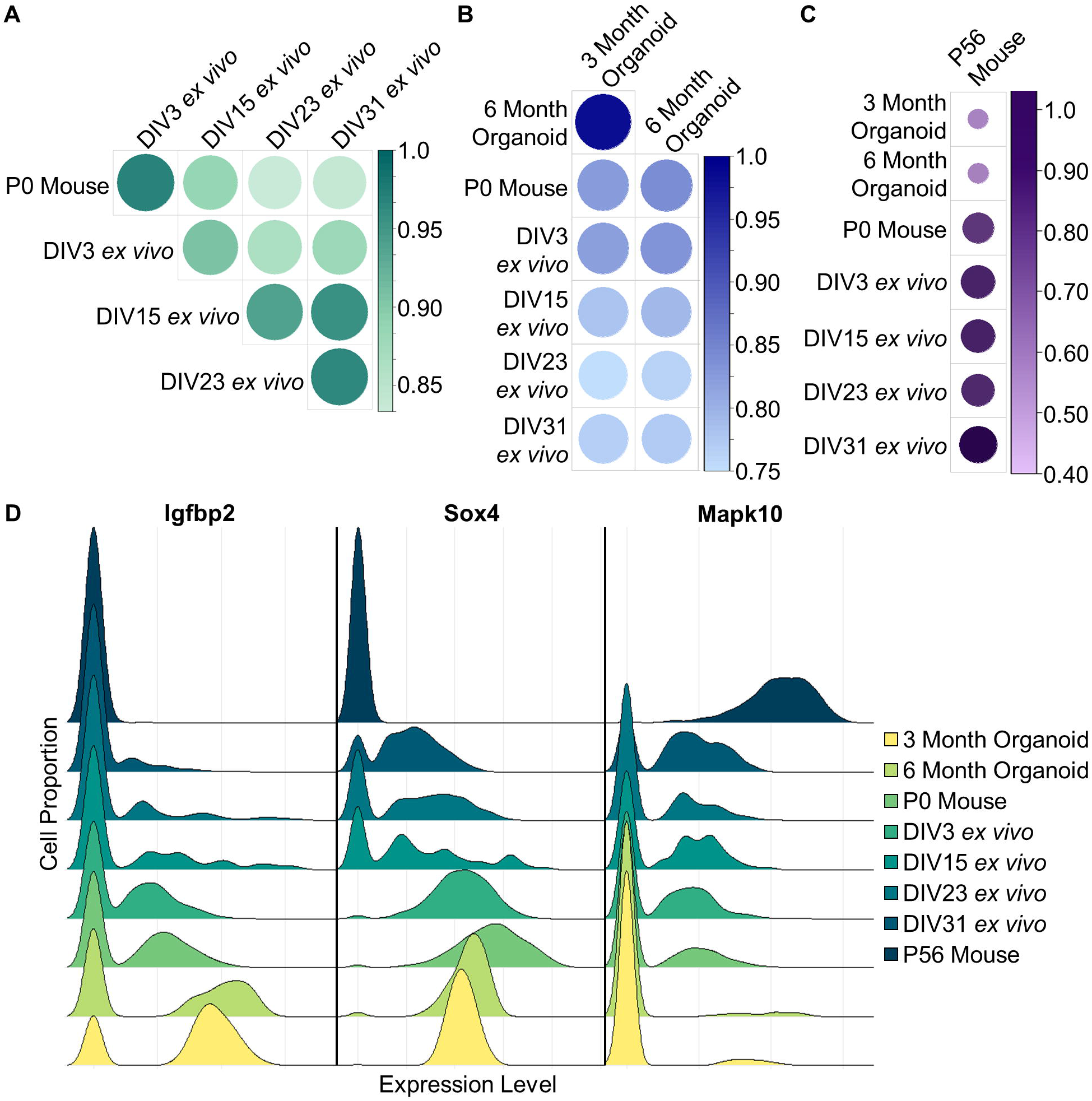
Cortical human organoids show developmental immaturity as compared to early postnatal mice (A) Correlogram displaying the correlation between all our developing mouse datasets (P0 mouse and DIV03-DIV31 *ex vivo*) for the average expression of all detected genes in the cortex excitatory neuron population. Color and circle size represent correlation value. (B) Same as (A), displaying the correlation of the two human cortical organoid datasets (3- and 6-month organoids) with each other and all the developing mouse model datasets for the average expression of all detected genes. (C) Same as (A), displaying the correlation of the adult (P56) mouse dataset with all other analyzed datasets for the average expression of each gene in only the cells that express that gene for the genes that are detected in all our analyzed models. (D) Ridgeline plot of the cortex excitatory neuron population for three representative genes, showing gene expressions for all models analyzed. Color represents model, from least to most developmentally mature.

We next looked at how the average gene expression for all genes from the two human cortical organoid time points correlate with each other, and with each of the timepoints in the developing mouse models (Figure 7B). The 3-month and 6-month organoids showed very high correlation with each other in each of the three major cell populations analyzed (0.95-0.98). When compared to the developing mouse datasets, both organoid timepoints are most highly correlated with P0 and DIV3 (0.77-0.84), and least highly correlated with DIV23 and DIV31 (0.72-0.78). Strikingly, the 6-month-old organoid is significantly more transcriptomically similar to the 3-month-old organoid than it is to the P0 mouse, demonstrating how relatively immature the human cortical organoid models are as compared to early postnatal mouse cultures.

We performed GO enrichment analysis on gene sets of the 500 genes with the largest average expression differences between the 3-month organoids and both the DIV3 and DIV31 datasets. For excitatory neurons, the genes with higher expression in the organoids as compared to both DIV3 and DIV31 were most strongly enriched for biological processes related to metabolism and mRNA processing, with less, but still significant, enrichment for processes related to differentiation and development. The genes with higher expression in the DIV3 and DIV31 datasets than the 3-month organoids were most strongly enriched for biological processes related to translation and cellular biosynthesis, with less, but still significant, enrichment for processes related to synaptic activity. The inhibitory neuron populations showed the same trends as the excitatory neurons, with the addition of mitotic processes being enriched for genes with higher expression in the organoids. Finally, the astrocyte populations had the same significant enrichments as the neuronal populations, with the absence of any significant enrichment for pathways related to development, differentiation, and synaptic activity.

Taking the list of all genes that are expressed in all four datasets (Organoid, P0, *ex vivo*, Adult), we investigated for each gene the average expression in all cells that have detectable expression of that gene. Using this data, we analyzed the correlation between the dissociated adult mouse cortices dataset with each of the developmental datasets (Figure 7C). In the excitatory neuron and inhibitory neuron populations, the adult dataset showed the highest average correlation with the DIV31 *ex vivo* dataset (0.58 and 0.53 respectively), and the lowest correlation with the 3-month human cortical organoid dataset (0.18 for both). The astrocyte population showed the same trend at a lesser magnitude, with a correlation of 0.38 to DIV31 *ex vivo* dataset and a correlation of 0.13 to 3-month human cortical organoid dataset. With all of this data, it is evident that there is a clear intrinsic progression of cortical development that spans species and developmental environment (Figure 7D).

### Unique expression profiles of disease genes highlight importance of disease model selection

Understanding which disease genes are expressed in each cortical model, and which are not, is critical for experimental design and model selection when investigating any gene of interest. Every epilepsy gene in our dataset had detectable expression in at least one of the models we analyzed, but as we have shown, this expression is not constant between models. Genes such as *Casr, Spr98*, and *Chrna2* only had detectable expression in the adult mouse, while genes such as *Map1b, Syngap1*, and *Sptan1* had expression in all models except for the adult mouse. Alternatively, *Cpa6, Nhlrc1*, and *Sik1* had no detectable expression in only the organoids. Examples of ASD genes with no expression in any of the models (*Hectd4, Kmt5b, Pax5*), expression in only the adult mouse (*Hras1, Mtap1a*), and no expression in only the adult mouse (*Hras, Kmt2e, Irf2bpl*) were also discovered. To show these details in their entirety for all the genes associated with neurological disorders we investigated, we have compiled the expression levels for each of the timepoints in the four models of cortical development analyzed (Table S15).

## Discussion

Here we describe longitudinal single cell transcriptomic analyses of mouse cortical and hippocampal cultures during early postnatal development. We interrogated cell-type and class-specific ontological trends, gene-specific expression patterns, and their involvement in genetic neurological diseases. These data provide a resource for studying cell population-specific longitudinal transcriptomic differences between and within different genetic model systems, species, and developmental maturities.

Neurodevelopmental disease modeling and therapy development face difficulties driven by brain complexity, the boundaries of *in vivo* and *in vitro* observations and perturbations, and the faithful structural and functional brain recapitulation, all of which hinder experimental design and analysis to a certain extent [35, 92–94]. These hurdles have even led to an attempt to forgo the physical brain entirely, and instead pursue large-scale *in silico* brain simulations [95–97]. A better understanding of the transcriptomic differences between different experimental models (e.g. human brain organoid vs. primary cultured neurons) allows one to better design disease modeling and/or screening experiments. Patient iPSC-derived brain organoids are now often used as model systems for neurological disease, brain development, and therapy testing. However, our data suggest that even after 6 months of development, cortical organoids are transcriptomically less mature than a newborn mouse (Figure 7C, D), which is itself more transcriptomically similar to a DIV31 cortical culture than to the organoid. This difference in intrinsic model maturity is crucial for their proper utilization when looking at genes associated with neurological disorders, including epilepsy, ASD, dystonia, and more (Table S9). For example, the epilepsy genes *Depdc5, Kcna2*, and *Adra2b* show little to no expression in the developing mouse primary cultures, while showing near ubiquitous expression in the adult mouse cortex (Table S15). In contrast, the NDD genes *Adnp, Map1b*, and *Prrt2* show high expression in human cortical organoids and low to no detectable expression in the developing primary cultures, nor in cortices of newborn or adult mice.

We identified disparate longitudinal transcriptomic patterns in excitatory neurons between genes associated with epilepsy and ASD, where epilepsy genes had an average peak expression at later (more mature network) timepoints, and ASD genes were generally more highly expressed at earlier timepoints in development (Figure 6A:B). This trend is weaker for ASD genes in inhibitory neurons but strengthened for epilepsy genes. Together these trends might be explained by the molecular functions of the genes associated with these diseases (Figure 6C) – many epilepsy genes are involved in pre- and postsynaptic processes and ion channel activities, which increase as the culture matures (Figure 7D) whereas, in contrast, many ASD genes are associated with neurodevelopmental processes including transcriptome regulation. These data suggest that when interrogating the functional or transcriptomic effects of most epilepsy genes and genes associated with functional maturity, utilizing mouse brain derived *ex vivo* networks are preferred in contrast to the more immature human brain organoids, as these genes have low or no detectable expression until later in network maturity. The opposite suggestion can be made for most ASD genes and other genes associated with brain development, where human organoid models might be better utilized. More broadly, our data demonstrate how well this ex vivo model recapitulates what is seen *in vivo*, and shows that for all diseases and biological processes analyzed, transcriptomic temporal trends are highly unique and require specific understanding of gene expression timing (Table S10).

Recent genetic discoveries and advances in computational approaches have provided new avenues for translational research such as transcriptomic reversal in neurological disorders[44, 98–101]. The genetic heterogeneity of NDDs partly underlies their pathophysiological and etiological variation. Accurately understanding both the developmental expression profile of the disease gene of interest and the effects of its loss (for loss of function mutations) on cell-type-specific transcriptomic profile are critical to assess the efficacy of pharmacological or genetic perturbations.

Together, the analyses presented here provide a wealth of transcriptomic data describing postnatal cortical and hippocampal *ex vivo* development, and its implication on different NDDs (Table S3, S12-15). This *ex vivo* model with other widely used cortical models, can be used as an agnostic tool to better understand the transcriptomic similarities and differences of distinct cell populations during neuronal network development, and provide guidance on experimental design when interrogating a specific gene, disease, or biological process.

## Acknowledgements

We thank Ryan Dhindsa for initial discussions on Single Cell Sequencing and for providing the P0 mouse dataset for our comparative analysis. We also thank Dr. Kristin Baldwin (CUMC) and Dr. Vincenzo Gennarino (CUMC) for valuable discussions regarding data analysis and experimental design.

## Author Contributions

Conceptualization, D.K.K., D.B.G., M.J.B.; Methodology, D.K.K., D.B.G., M.J.B; Validation, D.K.K.; Formal Analysis, D.K.K.; Investigation, D.K.K.; Resources, M.J.B, D.B.G.; Data Curation, D.K.K.; Writing – Original Draft, D.K.K.; Writing – Review & Editing, D.K.K., M.J.B.; Visualization, D.K.K., M.J.B; Supervision, M.J.B., D.B.G.

## Methods

### Mouse Husbandry

All data was generated and experiments were performed using inbred background C57BL/6J (000664 JAX stock) male mice. All mice were maintained in ventilated cages with controlled humidity at ∼60%, 12h:12h light:dark cycles and controlled temperature of 22–23°C. Mice had access to regular chow and water without restriction. Breeding cages were fed a high fat breeder chow. Mice were maintained and all procedures were performed within the Columbia University Institute of Comparative Medicine, which is fully accredited by the Association for Assessment and Accreditation of Laboratory Animal Care. All protocols were approved by the Columbia Institutional Animal Care and Use Committee.

### Primary Neuronal Culture

Prior to use, 48-well CytoView MEA plates (Axion BioSystems) and 24-well plates (Corning) with and without cover slips were coated overnight with 50 μg/mL poly-D-lysine (Sigma) in 0.1M borate buffer (pH 8.5). Before use, all plates were washed twice and allowed to dry at room temperature for at least 1h. Postnatal day 0 (P0) cortices and hippocampi acquired from 12 male mice from 3 separate litters were combined and dissociated using activated 20 U/mL Papain/DNase (Worthington) diluted in Hibernate A (Gibco) for 15 minutes at 37°C. Following manual trituration using a P1000 tip, cells were centrifuged at 300 x g for 5 min, and washed in 1X PBS. Cell pellets were suspended in NBA/B27 [consisting of Neurobasal-A (Life Technologies), 1X B27 (Life Technologies), 1X GlutaMax (Life Technologies), 1% HEPES, and 1% Penicillin/Streptomycin], supplemented with 1% fetal bovine serum (Gibco) and 5 μg/mL laminin. For the MEA plate, 50,000 cells were plated in a 40 μl convex droplet over the electrodes, incubated for 1h, then and additional 460 μl of media was added. 500,000 and 250,000 cells were plated in the 24-well plates for scRNA-seq and ICC respectively in a volume of 500 μl. The day after plating, media was removed and replaced with pre-warmed NBA/B27 without fetal bovine serum and laminin. Cultures were maintained at 37°C in 5% CO2. 50% of the medium was changed every other day with NBA/B27 starting on DIV3.

### Multi-electrode arrays (MEA)

Recordings were conducted every other day starting DIV3 prior to media change for 15 min per day using Axion BioSystems Maestro 768 channel amplifier and Axion Integrated Studios software (v2.4) at 37C in a carbogen mixture of 95% O2, 5% CO2. Each well is comprised of 16 electrodes on a 4 by 4 grid with each electrode capturing activity of all nearby neurons. A Butterworth band-pass filter (200–3000 Hz) and an adaptive threshold spike detector set at 7x the standard deviation (SD) of noise was used to record raw data and spike list files. Data were analyzed using the meaRtools package[102]. Spike list files were used to extract additional spike, burst, and network features. At least 5 spikes/min per electrode were required. Wells with fewer than four active electrodes for more than 30% of the total recording days were discarded. For burst detection, the maximum interval burst detection algorithm (Neuroexplorer software, Nex Technologies), which has been implemented into meaRtools, was used. As done in previous studies [47, 49, 103], we required that a burst consist of at least five spikes, lasting at least 50 ms, and that the maximum duration between two spikes within a burst was 0.05 s. Adjacent bursts were merged if the duration between them was <0.11 s.

### Immunocytochemistry

At DIVs 3, 9, 15, 23, and 31, cells were quickly washed with phosphate buffered saline (PBS) then fixed with 4% paraformaldehyde for 30 min at RT. Cells were washed 3x with PBS then incubated with blocking solution (5% normal donkey serum, 1% bovine serum albumin, 0.3% Triton X-100 in PBS) for 1 hour at RT. Cells were then incubated with primary antibodies diluted in blocking solution for 1.5 hours at RT, subsequently washed three times with 0.2% Triton X-100 in PBS, then incubated with the secondary antibodies diluted in blocking solution for 30 min at RT in the dark. The cells were then washed two times with 0.2% Triton X-100 in PBS, one time with PBS, and coverslips were mounted on microscopy slides with one drop of Prolong Antifade DAPI (Invitrogen) and allowed to cure in the dark overnight at RT, then imaged. Primary antibodies: Synapsin 1 (SYN) antibody (106 011, Synaptic Systems, 1:1000, mouse), Glial Fibrillary Acidic Protein (GFAP) antibody (Z0334, Agilent Dako, 1:2000, rabbit), MAP2 antibody (188 004, Synaptic Systems, 1:1000, guinea pig). Secondary antibodies: Donkey anti-Mouse IgG (H+L) Highly Cross-Adsorbed Secondary Antibody, Alexa Fluor 568 (A-10037, Invitrogen, 1:1000, donkey), Donkey anti-Rabbit IgG (H+L) Highly Cross-Adsorbed Secondary Antibody, Alexa Fluor 640 (A-32802, Invitrogen, 1:1000, rabbit), Goat anti-Guinea Pig IgG (H+L) Highly Cross-Adsorbed Secondary Antibody, Alexa Fluor 488 (A-11073, Invitrogen, 1:1000, goat). Imaging and post-processing was performed with an inverted Zeiss AxioObserver Z1 epifluorescent microscope equipped with an Axiocam 503 mono camera, and images acquired with the Zen 2 software.

### scRNA-seq analysis

For both cortical and hippocampal cultures, at DIVs 3,9,15,23, and 31 the cells from two wells were washed 2x with PBS, dissociated with activated 20 U/mL Papain, gently triturated with P1000 pipette, and combined. Cells were then collected and submitted for scRNA-seq on the 10X Chromium system at the Columbia Sulzberger Genome Center. scRNA-seq libraries were constructed with the 10X Chromium Single Cell 3’ Reagent Kits v2, and samples were sequenced on a NovaSeq 6000. Reads were aligned to the mm10 genome using the 10X CellRanger pipeline with default parameters to generate the feature-barcode matrix.

Seurat v4 was used to perform downstream QC and analyses on feature-barcode matrices[70–72]. DIV9 datasets for both cortex and hippocampus had mean reads per cell that were too high and mean genes per cell that were too low, so this timepoint was removed from downstream analysis. For remaining four timepoints, we removed all genes that were not detected in at least 4 cells. We further removed cells with fewer than 500 genes or more than 5000 genes detected and all cells with greater than 15% of reads mapping to mitochondrial genes. 5,804 and 6,774 cells remained for cortex and hippocampus respectively after filtering.

The filtered matrices were log-normalized and scaled to 10,000 transcripts per cell. We then identified the top 2,000 most variable genes per sample and harmonized gene expression across datasets before clustering. We next identified anchors between samples in each dataset using the Seurat FindIntegrationAnchors function with the default parameters and then used the IntegrateData function to compute and integrated expression matrix. Next, the ScaleData function was used on this matrix to regress out the number of UMIs and mitochondrial read percentage per cell using linear regression. We then used the RunPCA function to perform dimensionality reduction. We then computed a cellular distance matrix using the top 30 dimensions to subsequently generate a K-nearest neighbor graph (KNN). With this KNN, we used the FindClusters function with a resolution of 0.8 to implement the Louvain Clustering algorithm, and visualized the cells using UMAP. Differentially expressed genes per cluster were identified using the FindMarkers function. These genes were then used to annotate and merge clusters based on expression of known canonical marker genes from previous single-cell publications[48, 73–76]. Downstream expression analysis was performed on these annotated clusters.

### RNA extraction and RT-qPCR analysis

RNA from cortex and hippocampus derived ex vivo cultures was extracted from 3 biological replicates for three time points (DIV3, DIV15, DIV31) using RNeasy Plus Mini Kit (Qiagen). cDNA was synthesized using a SuperScript IV Reverse Transcriptase cDNA synthesis kit (Invitrogen) using 250ng per sample. Quantitative real-time PCR was performed on a QuantStudio 5 (Applied Biosystems) using TaqMan Universal PCR Master Mix (Applied Biosystems) with two technical replicates per reaction. Selected TaqMan primers were: *Mt1* (Mm00496660_g1, ThermoFisher, FAM-MGB); *Map1b* (Mm00485261_m1, ThermoFisher, FAM-MGB); *Cck* (Mm00446170_m1, ThermoFisher, FAM-MGB); *Gnb1* (Mm00515002_m1, ThermoFisher, FAM-MGB). All data were normalized to *Gnb1*. Average Ct for each qPCR reaction was calculated by averaging Ct values between technical replicates across biological replicates. ΔCt was calculated using the respective *Gnb1* Ct value as control. Relative gene expression changes over time (-ΔΔCt) were calculated by subtracting the geometric mean of the three DIV3 biological replicate ΔCt values for each gene tested in both brain regions from each calculated ΔCt value.

### Human organoid, P0, and Adult mouse dataset analysis

The human 3 month and 6 month cortical organoid datasets used for comparison were found in the Gene Expression Omnibus, accession number <u>GSE129519</u> [29]. Cells were randomly sampled to reduce cell number to an amount similar to the *ex vivo* dataset, and subsequently processed using the Seurat pipeline in the same manner as done with the ex vivo datasets described above. After filtering for number of genes and percent mitochondrial gene expression, 6209 cell remained. The P56 adult cortical dataset used for comparison was found in the Gene Expression Omnibus, accession number <u>GSE71585</u> [74]. This data was processed in the same manner as the generated ex vivo data as described above, with the only difference being we removed cells with fewer than 500 genes or more than 12500 genes, with 1772 cells remaining after filtering. The P0 cortical seurat object used for comparison was acquired from a member of our lab, and can be found in following publication [48]. This data was processed in the same manner as done with the *ex vivo* datasets described above, with the only differences being they removed cells with fewer than 1000 genes or more than 5000 genes, and they we removed all cells with greater than 8% of reads mapping to mitochondrial genes.

## Excel Table Titles

Table S3: Cortex and Hippocampus *ex vivo* gene expression data

Table S7: Gene expression changes for representative GO pathways

Table S8: Genes with stable *ex vivo* longitudinal expression

Table S9: Disease gene lists

Table S10: Disease gene expression trends

Table S11: Disease genes with no expression

Table S12: Human cortical organoid gene expression data

Table S13: New born postnatal day 0 mouse cortex gene expression data

Table S14: Adult postnatal day 56 mouse cortex gene expression data

Table S15: Disease gene expression data across all models

