## Supplemental Figures and Tables for "Single Cell transcriptional analysis of *ex vivo* models of cortical and hippocampal development identifies unique longitudinal trends"

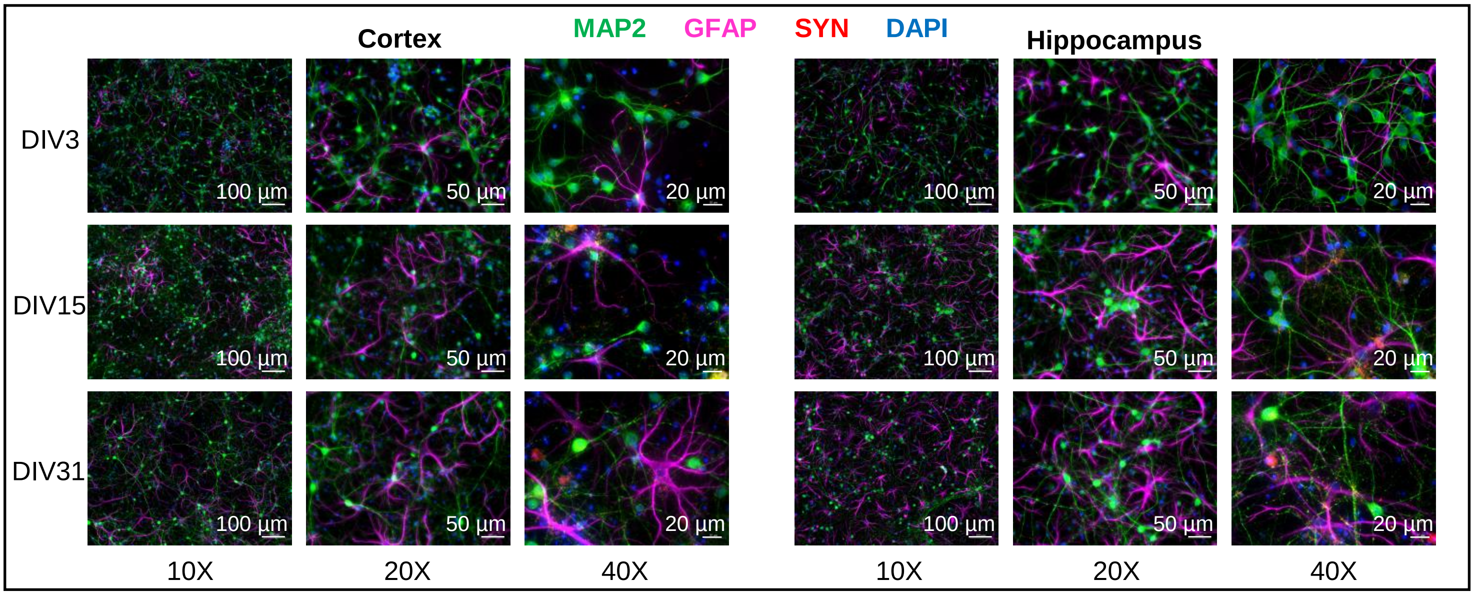


Figure S1. ICC images of cortex and hippocampus derived *ex vivo* cultures show proper network development

Immunocytochemistry of DIV3, DIV15, and DIV31 cortex and hippocampus derived cultures at 10X, 20X, and 40X magnification. Stained for neuronal (MAP2), glial (GFAP), and presynaptic (SYN) markers.

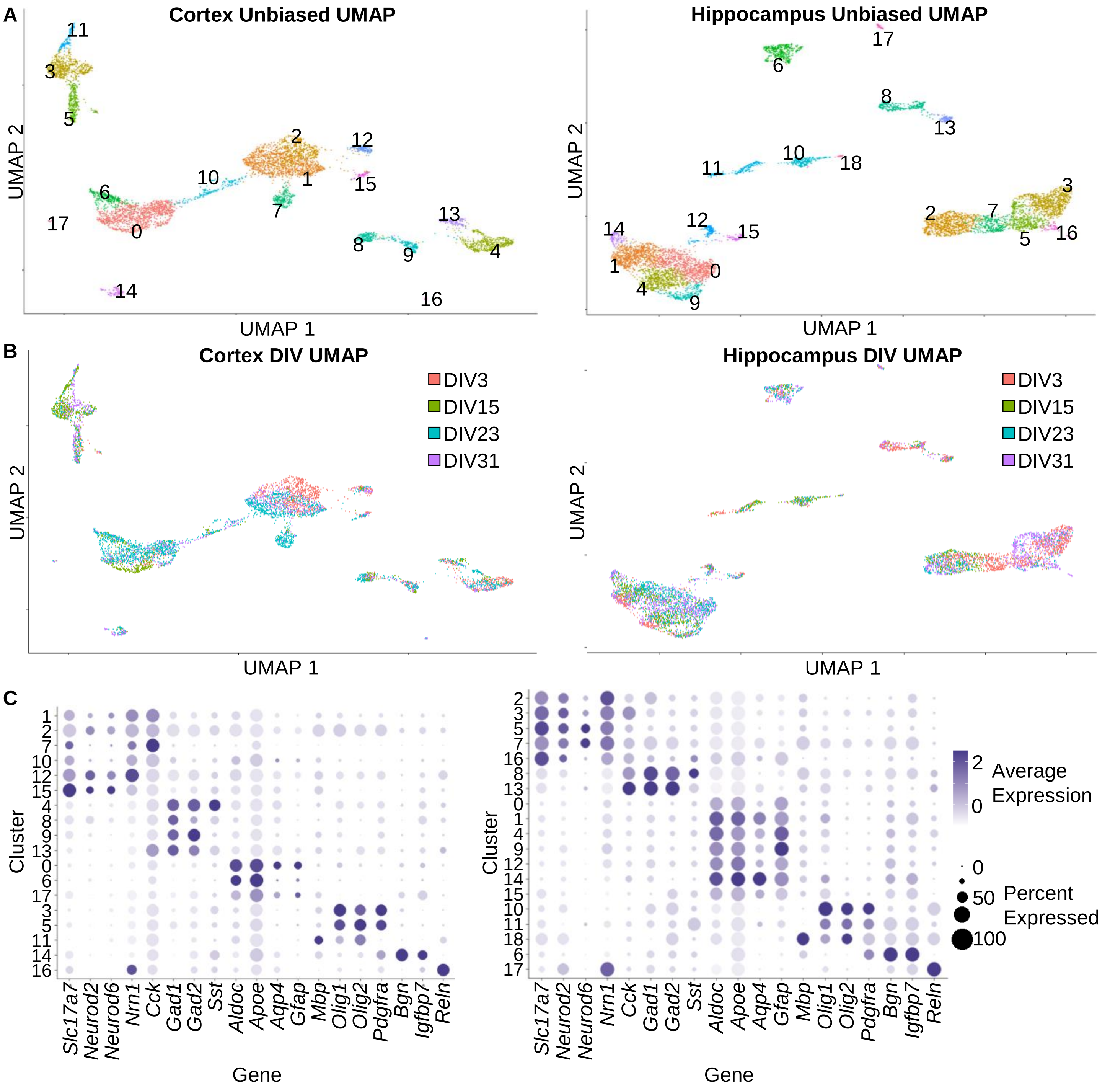


Figure S2. Unbiased single-cell RNA-sequencing of wildtype mouse cortex and hippocampus derived *ex vivo* cultures

(A) UMAP plot of scRNA-seq data from mouse cortex (left) and hippocampus (right) derived *ex vivo* cultures. Color represents unbiased cell clusters.

(B) Same as (A). Colors represent DIVs.

(C) Dot plots displaying the expression of select cell-type-specific marker genes (columns) in our cell clusters found in (A) for cortex (left) and hippocampus (right). Dot size represents percentage of cells expressing a marker gene, and color intensity represents relative average expression of a marker gene.

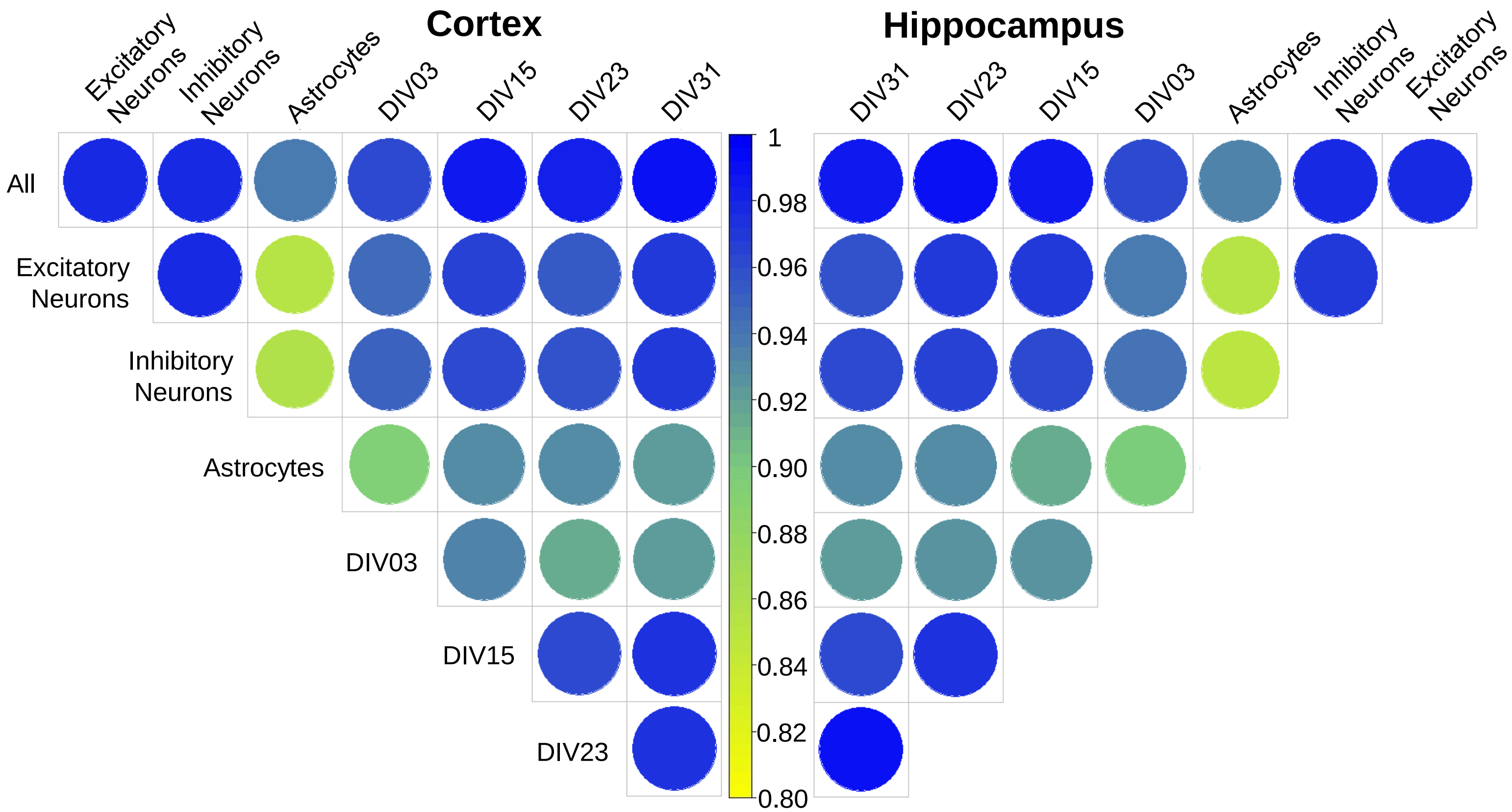


Figure S3. High gene expression correlation found between samples
Correlogram showing correlations of gene expression between major cell populations (Excitatory Neurons, Inhibitory Neurons, Astrocytes) and culture ages (DIV3, DIV15, DIV23, DIV31) for cortex (left) and hippocampus (right) derived ex vivo cultures. Colors represent gene expression correlation from high (blue) to low (yellow).

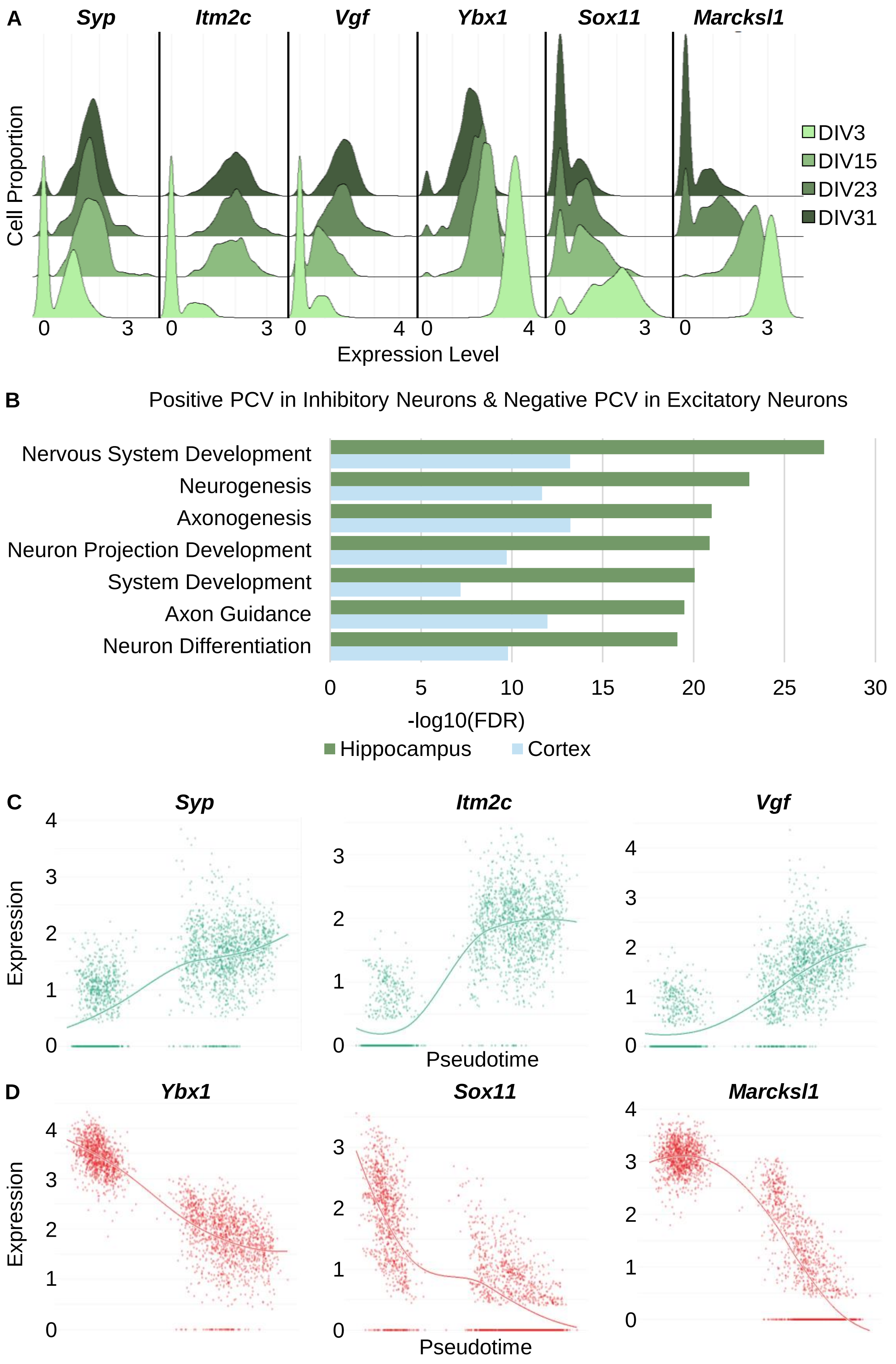

A
Syp
Itm2c
Vgf
Ybx1
Sox11
Marcksl1
Cell Proportion
DIV3
DIV15
DIV23
DIV31
Expression Level
B
Positive PCV in Inhibitory Neurons & Negative PCV in Excitatory Neurons
Nervous System Development
Neurogenesis
Axonogenesis
Neuron Projection Development
System Development
Axon Guidance
Neuron Differentiation
-log10(FDR)
Hippocampus
Cortex
C
Syp
Itm2c
Vgf
Expression
Pseudotime
D
Ybx1
Sox11
Marcksl1
Expression
Pseudotime

Figure S4. Hippocampus derived ex vivo cultures show similar trends as cortex derived cultures
(A) Ridgeline plot of the hippocampus excitatory neuron population for three representative genes with gene expression positively correlated with increasing culture age (left) and negatively correlating with increasing culture age (right). Color represents cell culture age.
(B) Gene ontology analysis of top 500 gene sets with the greatest difference in PCV where a gene has a positive PCV in the inhibitory neuron population and a negative PCV in the excitatory neuron population. Green bars represent the hippocampus derived ex vivo cultures and blue bars represent the cortex derived ex vivo cultures.
(C) Scatter plots comparing gene expression and pseudotime for the three genes in (Fig. 3B) that are positively correlated with culture age for each cell in the hippocampal excitatory neuron population. The line on each plot represents a smooth local regression of the data.
(D) Same as (B) for the three genes in (Fig. 3B) that are negatively correlated with culture age

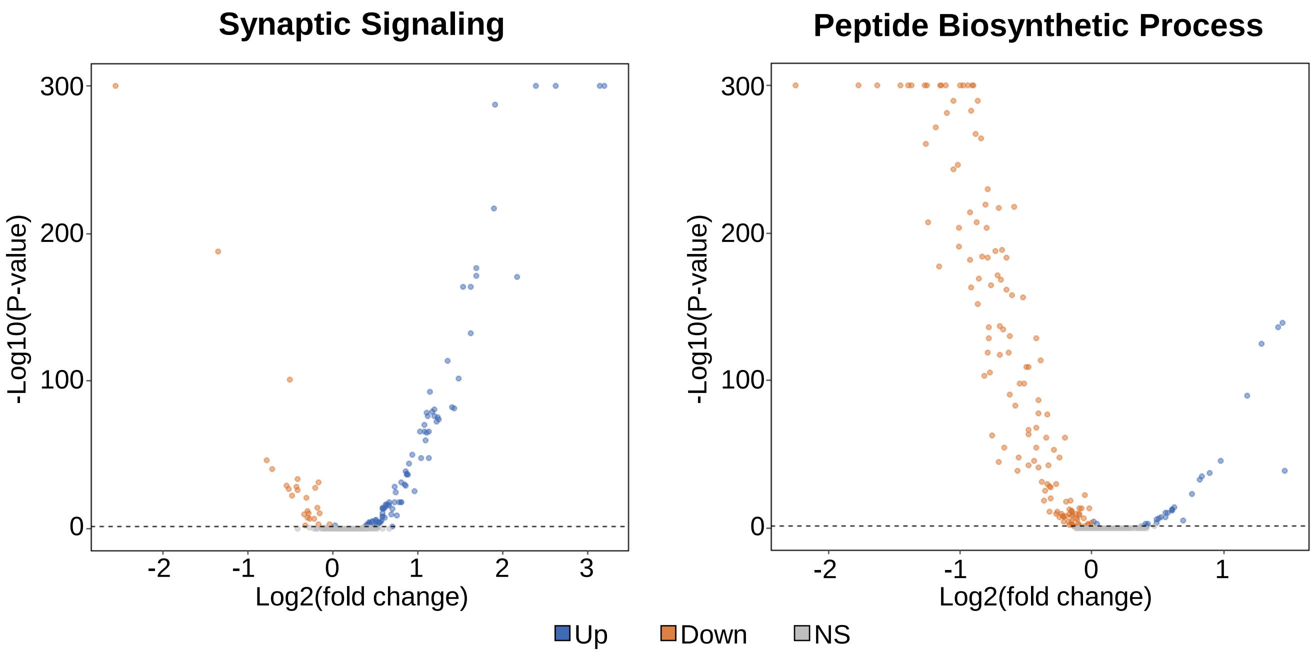


Figure S5. Gene expression changes between DIV3 and DIV31 for GO enriched biological pathways show strong directional trends

Volcano plots showing significant up and down regulation of gene expression from DIV3 expression to DIV31 expression for the cortex excitatory neuron population. Left plot is the 241 genes with detectable expression associated with the GO enriched biological pathway “Synaptic Signaling”. Right plot is the 313 genes with detectable expression associated with the GO enriched biological pathway “Peptide Biosynthetic Process”. Log2 fold change is represented on x-axis, and –log10 FDR adjusted p-value is represented on y-axis. Significant upregulation, significant downregulation, and no significance are represented with blue, orange, and grey datapoints respectively. P-value of 0 is represented with –log10(p-value) of 300.

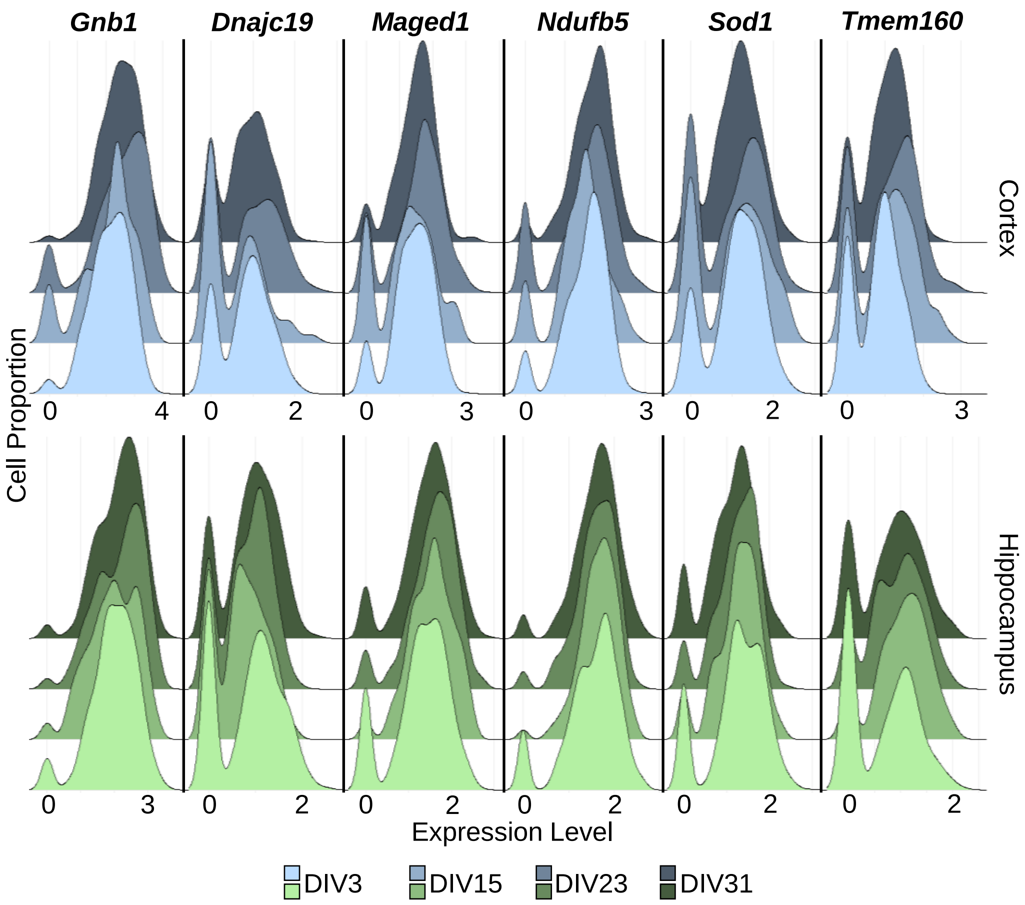


Figure S6. Housekeeping ridge plot of new genes
Ridgeline plots for six representative genes with stable gene expression through culture development for both the cortex derived ex vivo cultures(top) and the hippocampus derived ex vivo cultures(bottom). Color represents cell culture age.

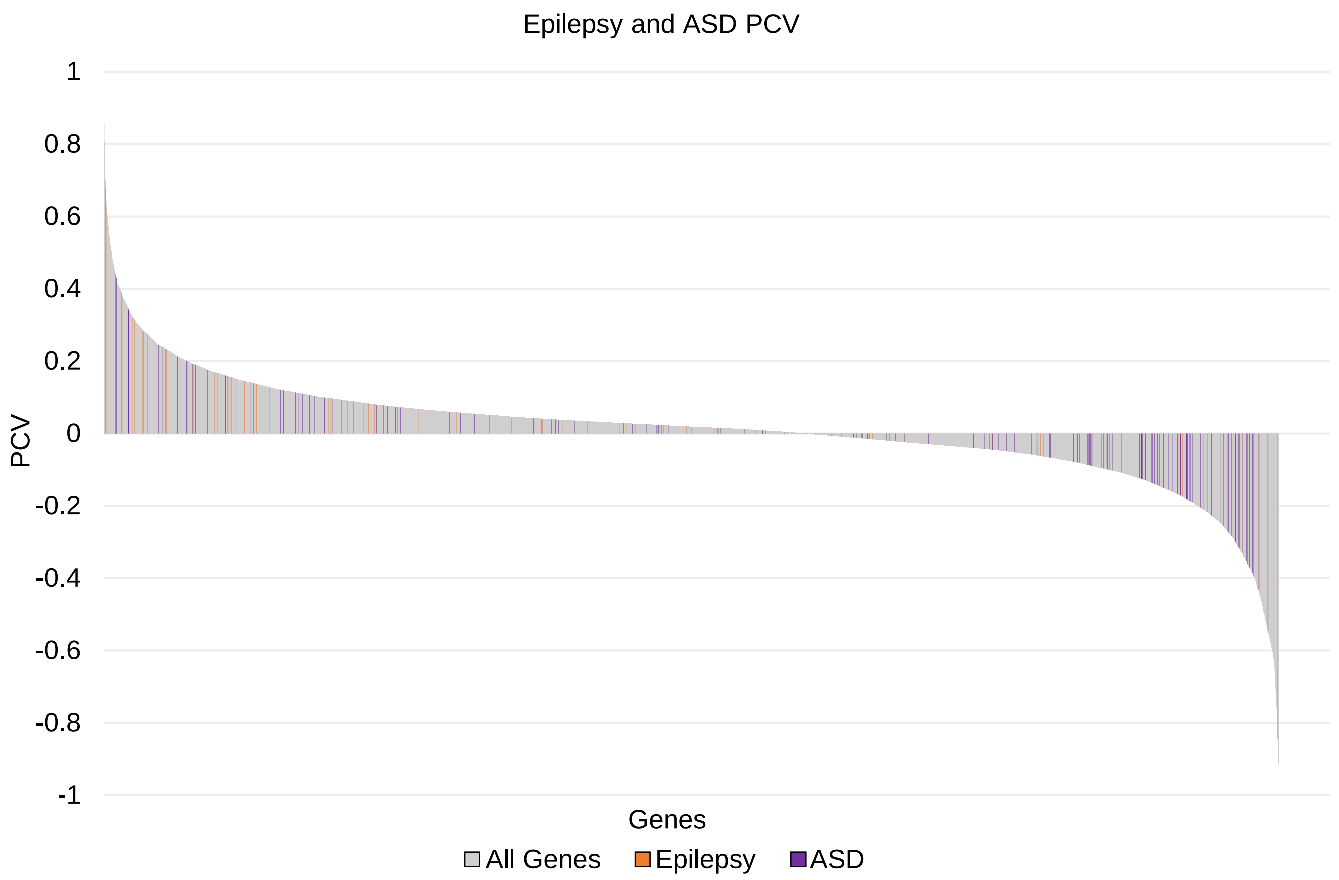


Figure S7. Epilepsy and ASD genes show distinct distribution of temporal expression patterns
Bar plot of all genes plotted from highest to lowest PCV along x-axis for cortex excitatory neurons. Autism Spectrum Disorder (ASD) genes and epilepsy genes are represented with purple and orange bars respectively.

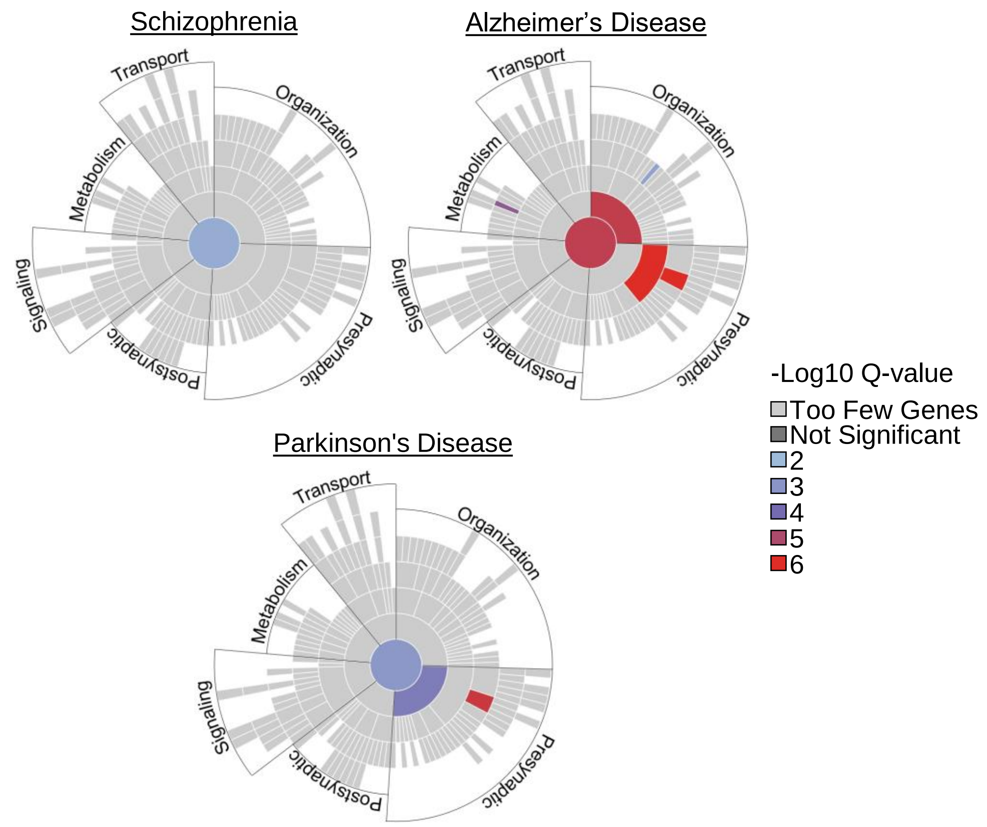


Figure S8. Disease genes show unique synaptic gene ontology profiles
SynGO plots for genes associated with schizophrenia (top-left), Alzheimer’s Disease (Top-right), Parkinson’s Disease (Bottom). Disease gene lists can be found in Table S8. Colors represent –log10 Q-values for enriched synaptic processes. Top-level synaptic processes are labeled. Dystonia, Obesity, and CAD had no significantly enriched synaptic processes.

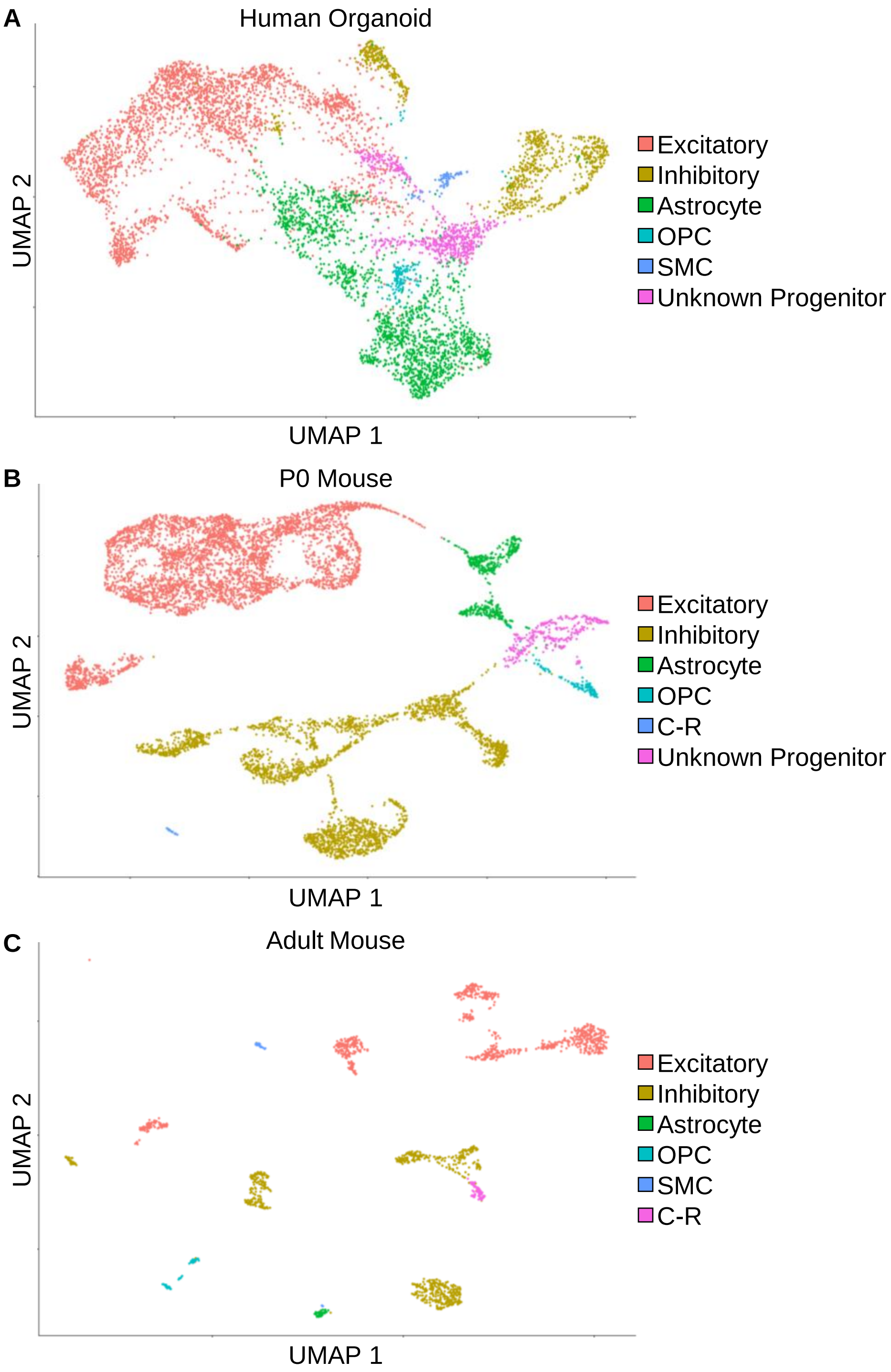


Figure S9. Distinct UMAP clustering found between models of cortical development
(A) UMAP plot of scRNA-seq data from 3 month and 6 month human cortical organoid cells. Color represents broad cell population clusters.
(B) UMAP plot of scRNA-seq data from newborn (P0) mouse cortical cells. Color represents broad cell population clusters.
(C) UMAP plot of scRNA-seq data from adult (P56) mouse cortical cells. Color represents broad cell population clusters.
Excitatory, Excitatory Neurons; Inhibitory, Inhibitory Neurons; OPC, Oligodendrocytes and Oligodendrocyte Precursor Cells (OPC); SMC, Smooth Muscle Cells (SMC); C-R, Cajal-Retzius Neurons.

| Brain Region | Days *in vitro* | Number of Cells | Mean reads per cell | Median genes per cell | Fraction reads in cells |
|---|---|---|---|---|---|
| Cortex | 3 | 1491 | 251183 | 2690 | 75.9 |
| Cortex | 9 | 6427 | 49206 | 280 | 49.4 |
| Cortex | 15 | 1366 | 272661 | 2543 | 52.8 |
| Cortex | 23 | 2365 | 159040 | 1906 | 52.8 |
| Cortex | 31 | 1066 | 321383 | 2577 | 55.4 |
| Hippocampus | 3 | 1985 | 191356 | 2194 | 69.3 |
| Hippocampus | 9 | 11757 | 32484 | 397 | 69.5 |
| Hippocampus | 15 | 2036 | 205095 | 3553 | 67.8 |
| Hippocampus | 23 | 2288 | 181713 | 3707 | 73.8 |
| Hippocampus | 31 | 2046 | 170239 | 2611 | 74.1 |

Table S1. *ex vivo* single cell 10x data

| | Cortex | | | | | Hippocampus | | | | |
|---|---|---|---|---|---|---|---|---|---|---|
| Cell Population | All | DIV03 | DIV15 | DIV23 | DIV31 | All | DIV03 | DIV15 | DIV23 | DIV31 |
| All | 5802 | 1351 | 1146 | 2328 | 977 | 6774 | 1874 | 1552 | 1489 | 1859 |
| Excitatory | 2242 | 814 | 93 | 964 | 371 | 2459 | 1044 | 306 | 338 | 771 |
| Inhibitory | 987 | 342 | 168 | 348 | 129 | 439 | 281 | 49 | 29 | 80 |
| Astrocyte | 1422 | 109 | 330 | 815 | 168 | 2973 | 369 | 844 | 924 | 836 |
| OPC | 1016 | 72 | 539 | 147 | 258 | 473 | 93 | 273 | 70 | 37 |
| SMC | 109 | 11 | 14 | 40 | 44 | 354 | 61 | 66 | 108 | 119 |
| Cajal-Retzius | 26 | 3 | 2 | 14 | 7 | 76 | 26 | 14 | 20 | 16 |

Table S2. *ex vivo* cell counts per population

| GO Biological Process | Fold Enrichment | -log10 (FDR) | GO Biological Process | Fold Enrichment | -log10 (FDR) |
|---|---|---|---|---|---|
| **Cortex Excitatory Neurons DIV3 > DIV31** | | | **Cortex Excitatory Neurons DIV31 > DIV3** | | |
| Nervous System Development | 2.59 | 16.66 | Regulation Of Transport | 3.02 | 23.26 |
| System Development | 1.84 | 11.95 | Localization | 1.85 | 16.10 |
| Regulation Of Macromolecule Biosynthetic Process | 1.91 | 11.92 | Regulation Of Ion Transport | 2.91 | 14.62 |
| Neurogenesis | 2.61 | 11.57 | Regulation Of Vesicle-Mediated Transport | 4.19 | 14.01 |
| Neuron Differentiation | 2.79 | 9.83 | Synaptic Signaling | 5.06 | 13.80 |
| Neuron Development | 2.90 | 8.65 | Ion Homeostasis | 3.33 | 10.82 |
| Neuron Projection Development | 3.12 | 8.38 | Regulation Of Synaptic Plasticity | 5.17 | 7.67 |
| **Cortex Inhibitory Neurons DIV3 > DIV31** | | | **Cortex Inhibitory Neurons DIV31 > DIV3** | | |
| Anatomical Structure Development | 1.55 | 3.63 | ATP Metabolic Process | 6.41 | 5.37 |
| Multicellular Organism Development | 1.57 | 3.59 | Carbohydrate Catabolic Process | 9.03 | 4.68 |
| System Development | 1.64 | 3.40 | Positive Regulation Of Transport | 2.43 | 3.97 |
| Developmental Process | 1.50 | 3.32 | Positive Regulation Of Cell Communication | 1.96 | 3.18 |
| Cell Differentiation | 1.54 | 1.75 | Positive Regulation Of Signaling | 1.95 | 3.17 |
| Nervous System Development | 1.75 | 1.64 | Positive Regulation Of Ion Transport | 2.59 | 3.16 |
| **Hippocampus Excitatory Neurons DIV3 > DIV31** | | | **Hippocampus Excitatory Neurons DIV31 > DIV3** | | |
| Anatomical Structure Development | 1.59 | 4.21 | ATP Metabolic Process | 5.94 | 3.91 |
| Multicellular Organism Development | 1.60 | 4.09 | Regulation Of Ion Transport | 2.27 | 3.79 |
| Developmental Process | 1.53 | 3.99 | Regulation Of Transport | 2.05 | 3.75 |
| System Development | 1.57 | 2.70 | Cellular Homeostasis | 2.49 | 2.94 |
| Regulation Of Multicellular Organismal Process | 1.61 | 1.30 | Ion Transport | 1.69 | 2.22 |
| Cellular Developmental Process | 1.52 | 1.27 | Establishment Of Localization | 1.51 | 1.92 |
| **Hippocampus Inhibitory Neurons DIV3 > DIV31** | | | **Hippocampus Inhibitory Neurons DIV31 > DIV3** | | |
| Anatomical Structure Development | 1.55 | 3.64 | Developmental Process | 1.47 | 3.00 |
| Developmental Process | 1.49 | 3.14 | Anatomical Structure Development | 1.46 | 2.59 |
| Multicellular Organism Development | 1.53 | 2.85 | Multicellular Organism Development | 1.45 | 2.07 |
| Animal Organ Development | 1.69 | 2.74 | ATP Metabolic Process | 4.44 | 2.05 |
| Regulation Of Multicellular Organismal Process | 1.65 | 1.80 | Cell Development | 1.74 | 1.47 |
| System Development | 1.50 | 1.77 | Positive Regulation Of Signaling | 1.74 | 1.37 |

Table S4. Gene ontology analysis shows differences between cell populations and brain region derived cultures

Gene ontology analysis of 500 genes with largest difference in average expression between DIV03 and DIV31 for cortex excitatory neurons, cortex inhibitory neurons, hippocampus excitatory neurons, and hippocampus inhibitory neurons. GO enriched biological processes and fold enrichment and –log10 values of FDR adjusted p-value (-log10(FDR)) are plotted in table.

| Gene | Cortex Log2 Fold Change: DIV3 vs DIV31 | Hippocampus Log2 Fold Change: DIV3 vs DIV31 | Cortex FDR adjusted p-value | Hippocampus FDR adjusted p-value |
|---|---|---|---|---|
| *Mt1* | 3.009860038 | 1.49679051 | 3.2091E-142 | 5.72147E-45 |
| *Map1b* | -2.451854406 | -1.353948639 | 0 | 2.5578E-132 |
| *Cck* | 2.86417 | -0.321519 | 2.70E-162 | 0.0000949361 |
| *Gnb1* | 0.108027463 | 0.120259293 | 1 | 1 |

Table S5. Genes validated with qPCR
Log2 fold change values between gene expression in DIV3 and DIV31 for both cortex and hippocampus derived ex vivo cultures and their respective FDR adjusted p-values. Mt1 has significant increased expression from DIV3 to DIV31 in both cortex and hippocampus. Map1b has significant decreased expression from DIV3 to DIV31 in both cortex and hippocampus. Cck has significant increased expression from DIV3 to DIV31 in cortex and significant decreased expression in hippocampus. Gnb1 has stable expression from DIV3 to DIV31 in both cortex and hippocampus. These genes were validated with qPCR, with Gnb1 used as reference gene.

| GO Biological Process | Fold Enrichment | -log10 (FDR) | GO Biological Process | Fold Enrichment | -log10 (FDR) |
|---|---|---|---|---|---|
| **Cortex Excitatory Neurons Low PCV** | | | **Cortex Excitatory Neurons High PCV** | | |
| Peptide Biosynthetic Process | 11.06 | 47.26 | Establishment Of Localization | 2.08 | 17.25 |
| Gene Expression | 3.05 | 27.00 | Ion Transport | 2.26 | 13.78 |
| Nervous System Development | 2.50 | 16.03 | Regulation Of Transport | 2.48 | 13.24 |
| Neurogenesis | 2.56 | 11.37 | Ion Homeostasis | 3.21 | 9.81 |
| Cellular Component Biogenesis | 2.19 | 11.49 | Synaptic Signaling | 3.98 | 8.06 |
| Neuron Differentiation | 2.68 | 9.05 | Regulation Of Vesicle-Mediated Transport | 3.14 | 6.91 |
| Regulation Of Cell Differentiation | 2.19 | 7.22 | Cell-Cell Signaling | 2.66 | 6.19 |
| **Cortex Inhibitory Neurons Low PCV** | | | **Cortex Inhibitory Neurons High PCV** | | |
| Translation | 15.02 | 71.40 | Localization | 2.12 | 28.95 |
| Cellular Macromolecule Biosynthetic Process | 4.36 | 33.28 | Ion Transport | 2.64 | 24.39 |
| Organic Substance Biosynthetic Process | 3.16 | 30.66 | Cation Transmembrane Transport | 5.13 | 17.52 |
| Cellular Metabolic Process | 1.94 | 30.47 | Regulation Of Trans-Synaptic Signaling | 4.53 | 15.11 |
| Biosynthetic Process | 3.09 | 29.91 | Regulation Of Synaptic Plasticity | 5.42 | 8.81 |
| Nervous System Development | 1.6 | 2.10 | Trans-Synaptic Signaling | 4.11 | 8.12 |
| **Hippocampus Excitatory Neurons Low PCV** | | | **Hippocampus Excitatory Neurons High PCV** | | |
| Nervous System Development | 3.00 | 26.47 | Establishment Of Localization | 1.94 | 12.50 |
| Neurogenesis | 3.15 | 20.01 | Localization | 1.74 | 11.78 |
| Axon Development | 6.09 | 18.82 | Transport | 1.93 | 11.71 |
| System Development | 2.01 | 17.84 | Ion Transport | 2.05 | 9.35 |
| Neuron Differentiation | 3.45 | 17.34 | Cation Transport | 3.02 | 7.52 |
| Axonogenesis | 5.97 | 16.51 | Ion Homeostasis | 2.73 | 6.29 |
| **Hippocampus Inhibitory Neurons Low PCV** | | | **Hippocampus Inhibitory Neurons High PCV** | | |
| Translation | 12.34 | 51.52 | Localization | 2.06 | 26.02 |
| Peptide Biosynthetic Process | 11.54 | 49.91 | Transport | 2.27 | 23.51 |
| Peptide Metabolic Process | 8.83 | 44.96 | Ion Transport | 2.38 | 17.50 |
| Gene Expression | 3.34 | 33.79 | Regulation Of Trans-Synaptic Signaling | 4.75 | 16.74 |
| Cellular Metabolic Process | 1.95 | 31.00 | Modulation Of Chemical Synaptic Transmission | 4.76 | 16.73 |
| Cellular Biosynthetic Process | 2.81 | 20.94 | Synaptic Signaling | 5.01 | 14.19 |

Table S6. Gene ontology analysis shows differences between cell populations and brain region derived cultures using PCV

Gene ontology analysis of 500 genes with largest and smallest PCVs for cortex excitatory neurons, cortex inhibitory neurons, hippocampus excitatory neurons, and hippocampus inhibitory neurons. GO enriched biological processes and their fold enrichment and –log10 values of FDR adjusted p-value (-log10(FDR)) are plotted in table.
